# Mitotic CDK4/6 activity sustains spindle checkpoint signalling to prevent mitotic slippage and genomic instability

**DOI:** 10.1101/2025.09.13.675969

**Authors:** Zhaoru Zhang, Yang Li, Maïté Leturcq, Yusanjiang Abula, Xinyue Wu, Yuu Kimata

## Abstract

The precise regulation of cell cycle entry and the maintenance of genome integrity are crucial for preventing tumorigenesis. Cyclin-dependent kinases 4 and 6 (CDK4/6) play pivotal roles in linking mitogenic signals to G1–S phase progression^1–3^, and their frequent deregulation in various cancers underscores their importance in driving cell proliferation and as therapeutic targets^4,5^. Despite this, the roles of CDK4/6 beyond the G1/S transition remain underexplored. Here, we uncover a previously unrecognised function of CDK4/6 in mitotic progression through regulation of the spindle assembly checkpoint (SAC)^6^. Using both cancer and non-transformed human cells, we show that acute CDK4/6 inhibition after G1/S transition leads to premature mitotic exit despite unattached kinetochores, resulting in chromosome missegregation and aneuploidy. Phosphoproteomic analyses and *in vitro* kinase assays reveal that CDK4 phosphorylates multiple sites on key SAC regulators, including the C-terminal tail of CENP-E, which is critical for BubR1 recruitment to kinetochores and SAC maintenance^7,8^. CDK4/6 inhibition reduces the phosphorylation of SAC components, attenuating checkpoint signalling and accelerating mitotic slippage. Notably, residual SAC activity persists, suggesting that CDK4/6 strengthens or stabilises SAC signalling rather than being essential for its activation. Thus, CDK4/6 functions extend beyond G1/S control to ensure mitotic fidelity, linking proliferating signals to genome stability and exposing potential vulnerabilities to anti-mitotic therapies in cancer.

## Introduction

Cyclin-dependent kinases 4 and 6 (CDK4/6), in association with D-type cyclins, are pivotal regulators of the G1 to S phase transition in the eukaryotic cell cycle, a critical checkpoint for cell proliferation^1–3^. CDK4/6 mediate this process by phosphorylating the retinoblastoma (Rb) protein, thereby releasing E2F transcription factors to activate genes essential for DNA replication and cell cycle progression^1,9^. Unlike canonical cell cycle CDKs, whose activities are intrinsically coupled with cell cycle phases, CDK4/6 activity is regulated by both extracellular mitogenic signals and internal cellular states, such as stress or developmental programs, primarily through the production of D-type cyclins and the levels of CDK inhibitors^1,10^. CDK4/6 also structurally diverge from classic cell cycle CDKs, forming a distinct subgroup with unique structural and functional characteristics^11^. Dysregulation of CDK4/6 is frequently observed in various cancers, including breast cancer and glioblastoma, where they drive oncogenesis through aberrant Rb pathway activation, underscoring their critical role as therapeutic targets. This has led to the development of several CDK4/6 inhibitors, including the FDA-approved drugs, palbociclib, ribociclib, and abemaciclib, which are widely used in treating certain malignancies, such as HR-positive, HER2-negative breast cancers^4,5^.

Emerging evidence indicates that CDK4/ extend their classical role beyond the G1/S transition. Recent studies employing single-cell analysis and CDK4/6-specific inhibitors have revealed their involvement in the G2/M transition, where inhibition of CDK4/6 can lead to cell cycle exit from the G2 phase^12,13^. Furthermore, CDK4/6 have been implicated in diverse non-cell cycle functions, including transcriptional and metabolic regulation^14,15^. In the context of cancer, CDK4/6 activity has been shown to influence immune system modulation and other pathways that contribute to tumour development and progression^16,17^. However, the full spectrum of CDK4/6 functions and the underlying molecular mechanisms remain largely unexplored.

The Spindle Assembly Checkpoint (SAC) is a conserved mitotic surveillance mechanism that ensures proper chromosome segregation and prevents aneuploidy. It monitors kinetochore-microtubule attachments and the tension across sister chromatids, delaying anaphase onset through inhibition of the anaphase-promoting complex/cyclosome (APC/C) until all sister kinetochores are bioriented^6,18,19^. Core SAC components include Mps1 and Aurora B kinases, the Mad1-Mad2 tetramer, and the mitotic checkpoint complex (MCC), complemented by additional regulators such as CENP-E and the ROD complex, which enhance checkpoint functionality^6,20,21^. While SAC activity is crucial for genome maintenance across all cells, its dynamics vary across cell types and during different stages of development, aging, and disease^22–24^. For instance, early embryos and germ cells often exhibit a higher rate of mitotic errors due to attenuated SAC activity^17,25–28^. In cancer, some cells develop an increased tolerance to chromosomal instability by upregulating SAC proteins such as Mps1, while others bypass mitotic arrest induced by anti-mitotic drugs through weakened SAC function, leading to mitotic slippage^29–31^. Intrinsic developmental programs and signalling pathways, such as the Ras-MAPK pathway, commonly activated by mitogens or hormones, have been implicated in regulating the expression levels and activity of key SAC proteins, thereby potentially influencing checkpoint sensitivity and robustness^25,32–35^. However, the mechanisms underlying SAC modulation and their significance remain poorly understood.

In this study, we identify a novel function of CDK4 in mitotic progression and chromosome segregation, extending its role beyond G1/S regulation. Our findings demonstrate that CDK4 regulates SAC and mitotic fidelity in cultured human cells. Specifically, inhibition of CDK4/6 after the G1/S transition or during SAC-mediated arrest results in mitotic exit with chromosome segregation failure, causing cells to accumulate in a 4N state. Mechanistically, CDK4 directly phosphorylates key SAC components, including CENP-E and CENP-F, highlighting its role in SAC modulation. This previously unrecognized function of CDK4 links mitogenic signalling intensity to cell division rates and mitotic accuracy. Furthermore, these findings suggest that CDK4 activity may confer tolerance to chromosomal instability and resistance to anti-mitotic therapies in cancer cells, presenting a potential avenue for therapeutic intervention.

## Results

### Acute CDK4/6 inhibition on cell cycle progression and Rb phosphorylation

Accumulating evidences suggest that CDK4/6 have roles beyond the G1/S transition. To investigate their phase-specific functions, we assessed CDK4/6 function in the cell cycle after the G1/S transition in cultured human cells. We chose a colorectal cancer cell line HCT116 as our primary experimental model. HCT116 is highly proliferative, with both CDK4 and CDK6 expressed and unmutated^36^, genetically relatively stable and highly amenable to genetic manipulations. In addition, we included two additional cancer cell lines: MCF-7, a breast cancer cell line commonly used to study CDK4/6 due to its frequent deregulation in breast cancer, and HeLa, a widely used cervical cancer cell, as well as an immortalised, untransformed retinal pigment epithelial cell line RPE-1 hTERT. To inhibit CDK4/6 activity, we utilised three clinically approved CDK4/6-specific inhibitors, palbociclib, abemaciclib and ribociclib (hereinafter, abbreviated into Palbo, Abema and Ribo, respectively), which are known for its specificity and has been extensively used in both research and clinical settings^37^.

We first assessed the ability of the distinct CDK4/6 inhibitors to induce G1 (or G0) cell cycle arrest and inhibit retinoblastoma (Rb) protein phosphorylation (pRb) in HCT116 cells. Asynchronously growing HCT116 cells were treated with varying concentrations of the CDK4/6 inhibitors for 24 hours. Cell cycle profiles were analysed by flow cytometry, and levels of phosphorylated Rb (pRb) were examined by Western blot (Extended Data Fig. 1a). All the three CDK4/6 inhibitors induced G1 arrest to varying degrees (Extended Data Fig. S1b, d). Palbociclib induced significant G1 accumulation and pRb reduction at concentrations of 0.15 μM and higher, reaching maximal effect at 5 μM (Extended Data Fig. 1b-e). Abemaciclib also showed G1 arrest at concentrations 0.15 μM or above, with maximal effect at 1 μM; however, at concentrations of 5 μM and higher, a slight increase in the G2/M population (cells with 4N DNA content) was observed (Extended Data Fig. 1d). Ribociclib also induced G1 arrest at concentrations of 1 μM and higher, with maximal effect at 5 μM. However, like abemaciclib, at 15 μM, the G2/M population was increased (Extended Data Fig. 1d). Correspondingly, all three CDK4/6 inhibitors reduced pRb levels in a dose-dependent manner in HCT116 cells (Extended Data Fig. 1c, e). Palbociclib and abemaciclib showed significant pRb reduction at concentrations of 0.15 μM and higher, with maximal inhibition observed at concentrations at 5 μM. Ribociclib required higher concentrations (1 μM and above) to achieve similar levels of pRb reduction, with maximal inhibition at 15 µM (Extended Data Fig. 1c, e).

We extended our analysis to MCF-7 and RPE-1 hTERT cell lines to assess the generality of palbociclib’s effects on G1/S progression. In MCF-7 cells, treatment with palbociclib at concentrations of 1 to 5 μM induced significant G1 accumulation after both 24 and 48 hours of incubation (Extended Data Fig. 1f). In RPE-1 hTERT cells, palbociclib concentrations of 0.5 μM and above caused clear G1 accumulation and pRb reduction, with maximal effect observed over 3 μM (Extended Data Fig. 1g, h). Thus, the other cell lines respond to palbociclib in a comparable kinetics to HCT116.

Based on these results, we concluded that a concentration of 5 μM palbociclib achieves maximal and specific CDK4/6 inhibition and primarily used this condition for subsequent experiments to examine CDK4/6 function in HCT116 cells. Notably, this concentration is comparable to the 3 µM used by Cornwell et al., 2023, which uncovered a role for CDK4/6 in maintaining CDK2 activity and pRb throughout interphase, beyond G1^13^.

### CDK4/6 inhibition after S phase initiation affects mitotic progression and fidelity

To examine the roles of CDK4/6 in specific phases of the cell cycle after the G1/S transition, we synchronised HCT116 cells at the G1/S boundary using a double thymidine block (see Methods). Following release, palbociclib was added at various time points to inhibit CDK4/6 and cell cycle progression was monitored (Fig. 1a, b). When introduced simultaneously with the release (T0), palbocilib caused a slight delay in S phase completion compared to control cells, likely due to the residual G1 cells transitioning into S phase being affected by CDK4/6 inhibition (Fig. 1a, "+Palbo after T0"). Additon of palbocilib at four hours post-release (T4), after cells had entered S phase, caused no significant effects on replication kinetics (Fig. 1a, "+Palbo after T4"). However, palbociclib-treated cells exhibited a delay in the accumulation of mitotic cells, indicated by 4N DNA content with positive staining for phosphorylated histone H3 at serine 10 (pH3) (Fig. 1a, T8-12). Extended analysis confirmed this delay in mitoitc accumulation and also showed delayed reaccumulation of G1 cells (2N DNA content with negative pH3 staining) (Fig. 1b, T9-15). In addition, a significantly larger G2 cell population (4N DNA content with no pH3 staining) persisted till the end of the time course (Fig. 1b, T15). These results indicate that acute CDK4/6 inhibition after the G1/S transition impacts the kinetics of mitotic progression, particularly the entry into mitosis. These results indicate that acute CDK4/6 inhibition after the G1/S transition affects the timing of mitotic progression, particularly entry into mitosis. The observation of a persisting 4N DNA cell population aligns with recent studies suggesting that CDK4/6 inhibition can lead to cell cycle exit during the G2 phase^12,13^.

**Figure 1.**
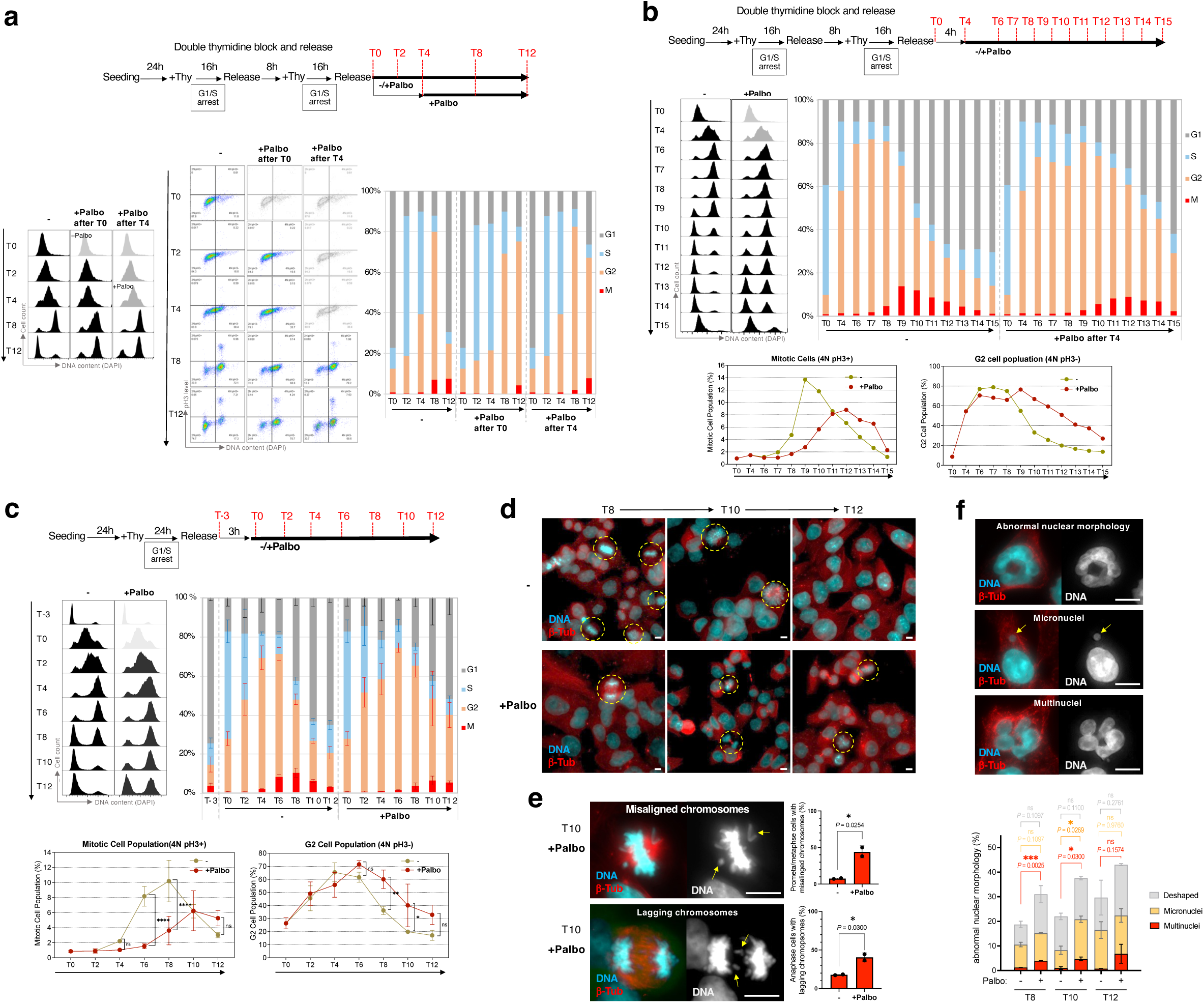
CDK4/6 inhibition after G1/S transition delays mitotic entry and compromises mitotic fidelity. **a, b,** HCT116 cells were synchronised at the G1/S boundary, treated with 5 µM Palbo or vehicle control at the indicated time points after release, and analysed by flow cytometry at the specified time points. **a**, Cell cycle profiles were monitored for 12 hours after Palbo addition at T0 or T4. DNA content and histone H3 serine 10 phosphorylation (pH3) signals were measured. **b,** Extended analysis with Palbo addition at T4. Proportions of mitotic cells (4N DNA content with pH3 signals) and G2 cells (4N DNA content without pH3 signals) were quantified over 15 hours. Palbo-treated cells displayed persistent G2 populations and delayed mitotic progression. **c,** Flow cytometry analysis of HCT116 cells treated with Palbo or vehicle control after synchronisation. Cell cycle profiles were monitored over 12 hours after Palbo addition. Means ± s.d. are shown (N=3). Statistical significance was assessed using two-way ANOVA. Palbo-treated cells showed delayed mitotic entry and increased retention in G2 phase. **d,** Representative immunofluorescence images of cells stained for DNA (Hoechst, cyan) and microtubules (β-tubulin, red) at T8, T10, and T12. Mitotic cells were shown in dotted yellow circles. Scale bars, 10 μm. **e, f,** Quantification of mitotic and post-mitotic abnormalities. **e,** Palbo-treated cells exhibited higher frequencies of chromosome congression defects (misaligned chromosomes) and lagging chromosomes during anaphase (yellow arrows). **f**, Palbo-treated interphase cells showed increased micronuclei (yellow arrows) and multinucleation. Quantification data are presented as means ± s.d. (N=2). For e, n = 127 and 134 (-); 90 and 89 (+Palbo) prometaphase and metaphase cells, and n = 81 and 115 (-); 85 and 81 (+Palbo) anaphase cells were analysed. For f, n = 500, 482 (-, T8); 492, 508 (+Palbo, T8), n = 493, 463 (-, T10); 475, 493 (+Palbo, T10), and n = 475, 521 (-, T12); 491, 486 (+Palbo, T12) interphase cells were analysed. Statistical significance was assessed using two-way ANOVA. In all statistical analyses presented in this study, significance is indicated as follows: ns: P ≥ 0.05, *: P < 0.05, **: P < 0.01, ***: P < 0.005, ****: P < 0.0001.

To characterise the effect of CDK4/6 inhibition on mitotic progression and chromosome segregation, we examined the morphology of cells that had passed through mitosis in the presence of palbociclib (Fig.1c-f). In our modified synchronisation protocol, cells were arrested at G1/S transition with single thymidine block and palbociclib was added three hours after release (Fig. 1c). As we observed with the double thymidine block, we observed slower accumulation of mitotic cells and higher retention of G2 cells upon palbociclib treatment, compared to untreated cells (Fig. 1c). Upon closer examination of mitotic cells, we found that palbociclib-treated cells displayed a higher frequency of chromosome congression defects and lagging chromosomes during anaphase (Fig. 1d, e). In addition, interphase cells with abnormal nuclear morphology, micronuclei, and multinucleation accumulated at later time points (T8-12, Fig. 1f). These findings suggest that CDK4/6 inhibition impairs mitotic fidelity, leading to chromosome missegregation and aneuploidy, highlighting a role for CDK4/6 in both proper mitotic progression and faithful chromosome segregation, in addition to its established function in promoting the G1/S transition.

### CDK4/6 inhibition causes premature mitotic exit

To understand how CDK4/6 inhibition affects mitosis and chromosome segregation, we first focused our analysis on the effect of CDK4/6 inhibition on early stages of mitosis. To this end, we blocked the metaphase-to-anaphase transition by activating the spindle assembly checkpoint (SAC) using nocodazole, a microtubule-depolymerising agent. HCT116 cells were synchronised at the G1/S boundary and then released into medium containing nocodazole, with or without CDK4/6 inhibition by palbociclib. The entry into and exit from mitosis were analysed by FACS. In the control condition, SAC effectively arrested cells in mitosis: Over 60% of cells showed 4N DNA content, with positive pH3 signals at 10–12 hours after release (T10-12, Fig. 2a, b, +Noc). However, in the presence of palbociclib, the accumulation of mitotic cells was significantly reduced: only approximately 20% of mitotic cells accumulated at 12 hours after release (T12, Fig. 2a, b, left, +Noc +Palbo). In addition, we observed gradual accumulation of cells in the G2-like status, lacking pH3 signals, but with 4N DNA content (Fig. 2a, b, right, +Noc +Palbo). Microscopic analysis revealed an increase in cells with multinuclei, common consequences of chromosome segregation failure (Fig. 2c). These results suggest that CDK4/6 inhibition induces premature mitotic exit or acceleration of mitotic slippage in mitotically arrested cells.

**Figure 2.**
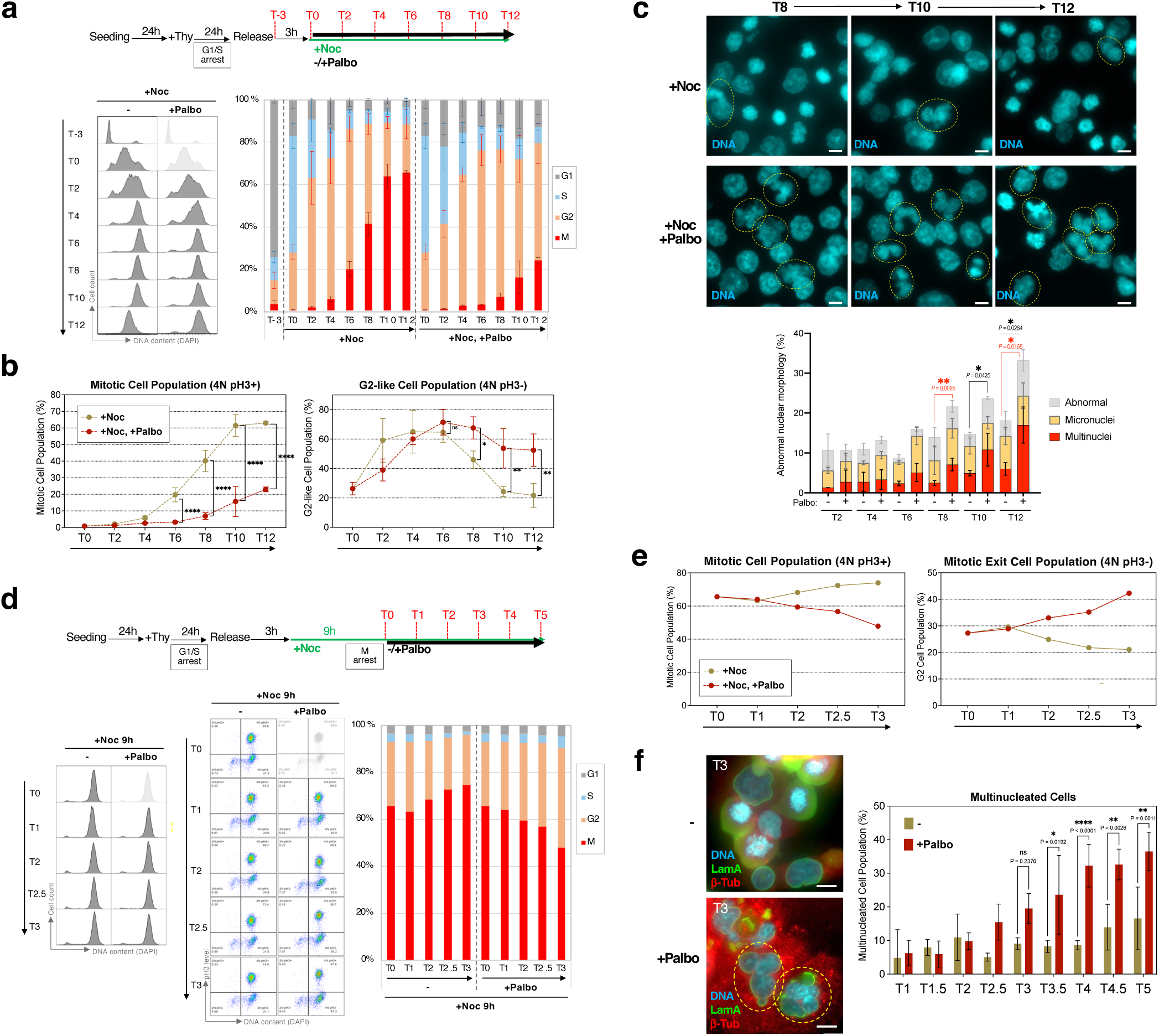
CDK4/6 inhibition promotes premature mitotic exit and mitotic slippage. **a, b,** HCT116 cells were synchronised at the G1/S boundary using a double thymidine block and released into nocodazole-containing medium to activate the spindle assembly checkpoint (SAC). Cells were treated with 5 µM palbociclib (Palbo) or vehicle control (+Noc) and harvested at the indicated time points for analysis by flow cytometry. **a,** Representative flow cytometry profiles showing DNA content (DAPI) and histone H3 serine 10 phosphorylation (pH3) at time points from T0 to T12. **b,** Quantification of mitotic cells (4N DNA content with positive pH3 signals, left) and G2-like cells (4N DNA content with negative pH3 signals, right). Palbo-treated cells showed reduced mitotic arrest and an increase in the G2-like population. Data are presented as means ± s.d. (N = 3). Statistical significance was assessed using two-way ANOVA. **c,** Representative immunofluorescence images of multinucleated interphase cells, indicative of mitotic exit following chromosome missegregation, stained for DNA (Hoechst, cyan), β-tubulin (red), and Lamin A/C (green). Yellow dashed circles indicate multinucleated cells. Scale bars, 10 μm. **d–f,** CDK4/6 inhibition in nocodazole-arrested mitotic cells induces mitotic slippage. **d,** Schematic of the experimental design and flowcytometry analysis: Cells were synchronised at G1/S, released into nocodazole-containing medium, and incubated for 9 hours to arrest in prometaphase. Palbo was added, and samples were collected at 0, 1, 1.5, 2, 2.5, and 3 hours after addition. **e,** Flow cytometry analysis of mitotic (4N pH3+) and G2-like (4N pH3−) cell populations over time. Data are presented as means ± s.d. (N = 3). **f,** ight: Multinucleated cells were quantified and presented as means ± s.d. (N = 3). Statistical significance was assessed using two-way ANOVA. The sample size (n) for each condition and time point across three biological replicates is as follows: replicate 1 (-: 123, 105, 102, 109, 109, 118, 142, 106, 117; +Palbo: 109, 113, 112, 114, 106, 101, 128, 124, 129), replicate 2 (-: 172, 93, 97, 108, 105, 126, 113, 93, 122; +Palbo: 123, 132, 116, 125, 132, 143, 135, 113, 123), replicate 3 (-: 192, 187, 190, 132, 195, 108, 184, 191, 175; +Palbo: 130, 309, 187, 121, 165, 162, 144, 128, 173) at T1 to T5.

To determine whether these mitotic phenotypes were caused by CDK4/6 inhibition during mitosis, or as the consequence of its inhibition prior to mitosis, we first arrested cells in prometaphase by treating them with nocodazole for 9 hours (over 60% of cells were mitotic) and then added palbociclib (Fig. 2d-f). In the control condition (“+Noc 9h, -”), cells remained arrested in mitosis, as evidenced by sustained high pH3 signals (Fig. 2d and 2e, left panel). However, upon addition of palbociclib, some cells prematurely exited mitosis, indicated by a decrease in pH3-positive cells and the simultaneous increase of pH3-negative cells (Fig. 2d and 2e, right panel). Concomitantly, multinucleated interphase cells also increased significantly (Fig. 2f). This suggests that CDK4/6 activity during mitosis is required for the maintenance of SAC-dependent mitotic arrest to prevent chromosome missegregation.

To confirm that this phenotype observed with palbociclib is specifically due to CDK4/6 inhibition, we tested the effects of other clinically used CDK4/6 inhibitors, abemaciclib and ribociclib, along with PD0325901, a specific inhibitor to MEK (MEKi). MEK, acting through the MAPK/ERK signalling pathway, influences CDK4/6 activity by modulating the expression and stability of Cyclin D^38,39^. HCT116 cells were arrested in mitosis by the incubation with nocodazole and then treated with increasing concentrations of the inhibitors, and the effects on mitotic exit were assessed by flow cytometry and Western blotting for Cyclin B1 (CycB1) (Extended Data Fig. 2a). Like palbociclib, abemaciclib induced premature mitotic exit, however, at lower concentrations than palboclibib: 0.5 μM and above. Ribociclib required higher concentrations (15 μM) to induce premature mitotic exit, which is expected given its higher inhibitory concentration for CDK4/6 than the other inhibitors^37^ (Extended Data Fig. 2b). MEKi did not affect SAC-dependent mitotic arrest at any of the concentrations tested (Extended Data Fig. 2b). Western blot analysis demonstrated that these effects on cell cycle progression correlated well with more rapid degradation of CycB1 as well as pRb reduction, confirming effective CDK4/6 inhibition (Extended Data Fig. 2c, d). Comparision of the effects of varying concentrations on premature mitotic exit showed similar kinetics between the three CDK4/6 inhibitors, but not to MEKi (Extended Data Fig. 2e). These data indicate that the induction of premature mitotic exit is specifically due to CDK4/6 inhibition and that the potency of the inhibitors corresponds to their ability to trigger this phenotype.

### CDK4/6 inhibition induces premature mitotic exit across multiple cell lines

To examine whether these effects of CDK4/6 inhibition is specific to HCT116 cell lines or can be extendable to other cells, we extended our analysis to other cancer cell lines: MCF-7 and HeLa cells, as well as untransformed RPE-1 hTERT cells. Since RPE-1 cells cannot be effectively synchronised using thymidine block^40^, we attempted to induce mitotic arrest by directly treating asynchronously growing cells with nocodazole without synchronisation. After 24 hours of nocodazole treatment, HeLa and RPE-1 cells accumulated in M phase (4N DNA content with pH3 signals) reaching over 60%. In contrast, HCT116 and MCF7 cells showed less mitotic population, reaching maximal G2/M populations of approximately 40% (Extended Data Fig. 3a, b).

We then treated these nocodazole-arrested cells with varying concentrations of palbociclib. In this assay, HCT116 cells showed accelerated mitotic slippage at 0.5 μM or more palbociclib (Extended Data Fig. 3a), as we observed with thymidine block (Fig. 2). In MCF7 cells, palbociclib also induced mitotic exit at concentrations of 1 μM and above (Extended Data Fig. 3b). In HeLa cells and untransformed RPE1 hTERT cells, palbociclib also induced mitotic exit, yet at higher concentrations, at 5 μM and above (Extended Data Fig. 3a, b). These findings indicate that the induction of premature mitotic exit by CDK4/6 inhibition is not cell-type-specific and is not due to thymidine pre-treatment, while different cells exhibit varying levels of sensitivity to palbociclib.

### Premature mitotic exit induced by CDK4/6 inhibition requires cyclin degradation

Our results demonstrate that palbociclib induces premature mitotic exit, causing chromosome missegregation in transformed and untransformed human cells. We explored the molecular mechanisms by which CDK4/6 inhibition causes premature mitotic exit. Mitotic exit necessitates the inactivation of CDK1 through the degradation of mitotic cyclins including CycB1 via the ubiquitin–proteasome pathway^41,42^. To determine whether cyclin degradation mediates the mitotic exit induced by CDK4/6 inhibition, we inhibited the 26S proteasome activity using MG132 (Fig. 3a). Cells were arrested in mitosis by nocodazole and mitotic exit was then induced by addition of palbociclib or the CDK1-specific inhibitor RO-3306, with or without co-treatment of MG132. Proteasome inhibition effectively blocked mitotic exit triggered by palbociclib, as evidenced by the retention of cells with positive pH3 signals (Fig. 3a, middle and bottom left). Immunoblot analysis showed that MG132 also caused the retention of high levels of CycB1, as well as phosphorylated forms of the APC/C subunit Cdc27, known to be phosphorylated by CDK1^43^, in cells treated with palbociclib (Fig. 3a, bottom right). Similarly, proteasome inhibition inhibited premature mitotic exit induced by abemaciclib (data not shown). In contrast, MG132 did not prevent mitotic exit nor Cdc27 dephosphorylation triggered by direct CDK1 inhibition (Fig. 3a, "+CDK1i"). These results indicate CDK4/6 inhibition induces premature mitotic exit through proteasome-dependent degradation of mitotic cyclins, rather than direct inhibition of CDK1 kinase activity.

**Figure 3.**
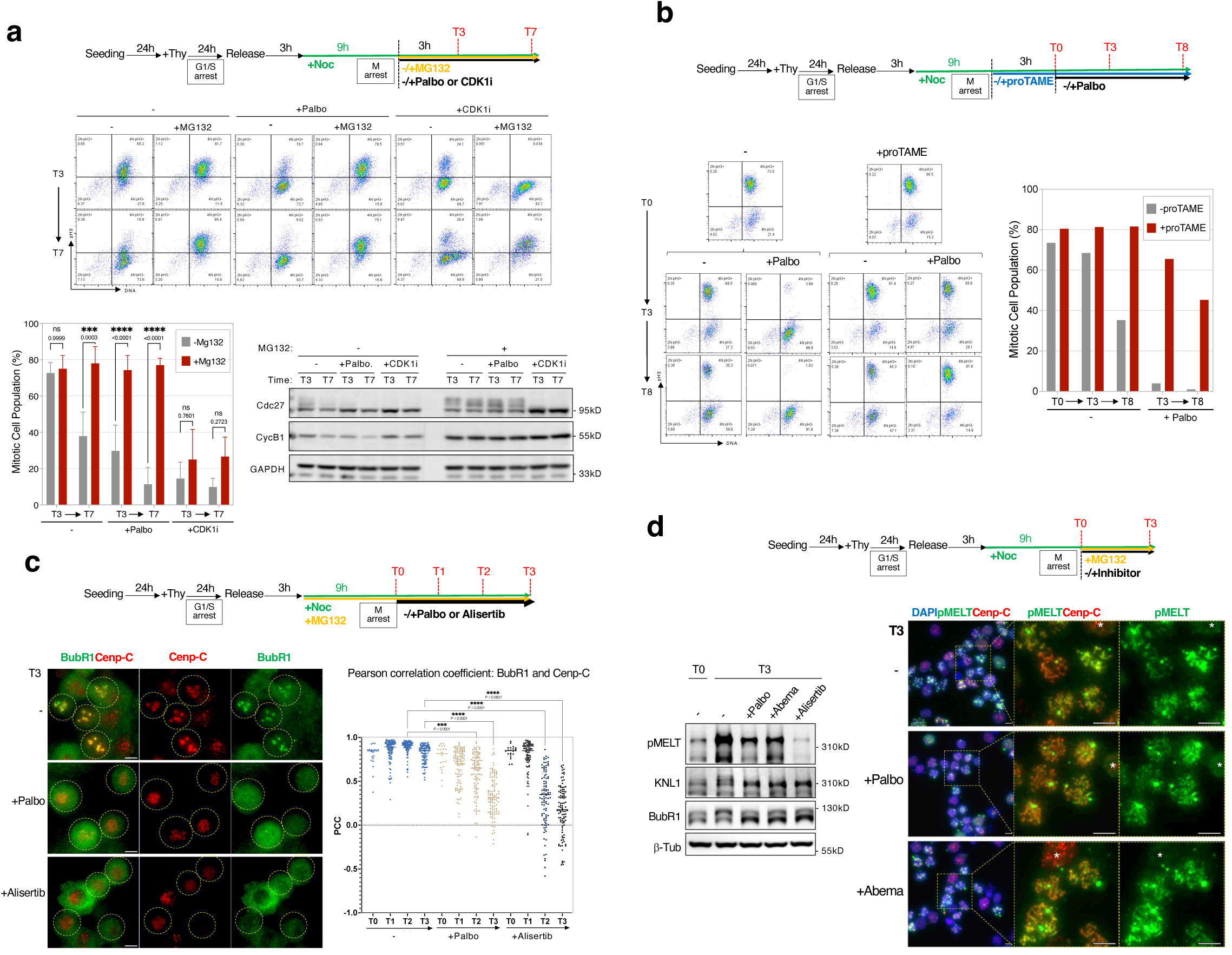
CDK4/6 inhibition leads to premature mitotic exit through spindle assembly checkpoint inactivation. **a,** HCT116 cells were synchronised in prometaphase using thymidine-nocodazole treatment, followed by 9 hours of nocodazole incubation. Cells were treated with Palbo or a CDK1-specific inhibitor, RO-3306 (CDK1i, 10 µM), with or without MG132. Samples were collected at 3 and 7 hours after treatment for flow cytometry and immunoblot analysis. Mitotic exit (decrease in pH3-positive cells) induced by Palbo, but not by CDK1i, was effectively blocked by proteasome inhibition. Data are means ± s.d. (N = 3); two-way ANOVA. Western blot analysis shows retention of highly phosphorylated Cdc27 and CycB1 in MG132-treated cells. **b,** Cells were synchronised in prometaphase as above and treated with Palbo or vehicle control in the presence or absence of the APC/C inhibitor, proTAME (25 µM). Flow cytometry analysis revealed that proTAME blocked mitotic exit induced by CDK4/6 inhibition, as evidenced by retention of pH3-positive cells with high CycB1 levels. **c,** HCT116 cells were arrested in mitosis using nocodazole and co-treated with MG132 to prevent mitotic exit. BubR1 and centromere marker CENP-C were visualised by immunofluorescence. BubR1 kinetochore localisation decreased over time upon Palbo treatment, while it remained stable in control conditions. Alisertib (1 µM) served as a positive control for SAC inactivation. Scale bar: 10 µm. Pearson correlation coefficients (PCC) quantify BubR1/CENP-C co-localisation; one-way ANOVA. n = 20, 100, 100, 100 (-); 20, 99, 100, 100 (+Palbo); 21, 100, 98, 100 (+Alisertib), at T0, T1, T2, T3 in a single experient. **d,** CDK4/6 inhibition reduces KNL1 MELT phosphorylation (pMELT) and BubR1 phosphorylation. Left: Immunoblots show decreased pMELT and BubR1 phosphorylation in cells treated with Palbo, Abema (5 µM), or alisertib in the presence of MG132, correlating with SAC inactivation. Right: Immunofluorescence shows partial reduction of kinetochore pMELT signals after Palbo or Abema treatment. Scale bar: 10 µm.

Mitotic cyclin degradation is initiated by the APC/C through ubiquitination^44,45^. To test if APC/C activity is required for the mitotic exit induced by CDK4/6 inhibition, we inhibited APC/C using proTAME, a specific APC/C inhibitor^46^. Similar to the data with MG132, preincubation with proTAME effectively inhibited cyclin degradation and prevented mitotic exit upon CDK4 inhibition, as shown by the retention of cells with high CycB1 and pH3 signals (Fig. 3b). These results indicate that CDK4/6 inhibition causes premature APC/C activation, leading to cyclin degradation and mitotic exit.

### CDK4/6 inhibition attenuates the spindle assembly checkpoint

APC/C activity is regulated by its co-activators CDC20 and CDH1 (also known as Fzr1): APC/C^CDC20^ is active from prophase until metaphase while APC/C^CDH1^ operates during late mitosis and G1 phase^44,45^. APC/C^CDC20^ activation is regulated by APC/C core subunit phosphorylation as well as the SAC, whereas APC/C^CDH1^ is inhibited by phosphorylation of CDH1 by CDK1 as well as CDK4^43,44,47,48^. It may be possible that CDK4/6 inhibition may induce cyclin degradation due to untimely APC/C^CDH1^ activation through removal inhibitory phosphorylation of CDH1. To test whether untimely CDH1 activation contributes to premature mitotic exit upon CDK4/6 inhibition, we utilised the AID2 system, developed in Kanemaki’s lab, to induce rapid depletion of CDH1^49^. We generated HCT116 cell lines where a plant-specific SCF substrate receptor TIR1 from *Oryza sativa* (OsTIR1) was ubiquitously expressed and the endogenous CDH1 gene was replaced with mini-AID (mAID)–mCherry-tagged CDH1 (Extended Data Fig. 4a). In these cells, upon addition of 5-phenylindole-3-acetic acid (5-Ph-IAA) to the media^49^, CDH1 was rapidly degraded, reaching below 20% of the original level after 3 hours (T0, Extended Data Fig. 4a). However, mitotic exit, as well as CycB1 degradation, still occurred upon CDK4 inhibition at comparable kinetics to control (Extended Data Fig. 4a). Thus, APC/C^CDH1^ activity is unlikely to contribute to the premature mitotic exit induced by CDK4/6 inhibition.

The SAC prevents APC/C^CDC20^ activation until all chromosomes have established proper bipolar attachments to the mitotic spindle and are aligned at the metaphase plate^18,19^. We next examined whether CDK4/6 inhibition compromises SAC function, potentially causing premature APC/C^CDC20^ activation. During SAC activation, core checkpoint components including BubR1 accumulate at unattached kinetochores, where the mitotic checkpoint complex (MCC) is formed to inhibit APC/C^CDC206,20,52^. To assess SAC integrity following CDK4/6 inhibition, we monitored BubR1 kinetochore localisation (Fig. 3c). Cells were arrested in mitosis using nocodazole and co-treated with MG132 to prevent mitotic exit. In control conditions, BubR1 remained highy accumulated with kinetochores, co-localising with the centromere marker CENP-C throughout the entire time course^18,21^ (Fig. 3c, "-"). By contrast, when cells were treated with palbociclib, BubR1 gradually dissociated from kinetochores ("+Palbo"), although this dissociation was slower than that induced by the Aurora A kinase inhibitor alisertib—previously shown to drive SAC inactivation^50^ ("+Alisertib").

Notably, alisertib treatment rapidly reduced pH3 signals on chromosomes, despite their highly condensed mitotic morphology, reflecting the inhibition of Aurora kinases, which have been shown to phosphorylate this site^51^ (Extended Data Fig. 4b). In contrast, palbociclib treatment did not alter pH3 levels, indicating that, under these conditions, CDK4/6 inhibition does not interfere with Aurora kinase activity (Extended Data Fig. 4b). Together, these observations suggest that CDK4/6 inhibition leads to premature SAC silencing and may allow cyclin degradation despite the presence of unattached kinetochores. This effect appears to occur in parallel with or downstream of Aurora kinases.

The phosphorylation of KNL1 and BubR1 are key steps in SAC signalling. Mps1 kinase phosphorylates the MELT repeats of KNL1 (pMELT), facilitating the recruitment of the Bub1-Bub3-BubR1 complex to kinetochores^52–55^. Additionally, BubR1 is phosphorylated by multiple kinases, including Plk1, CDK1 and Mps1, during mitosis, which is crucial for its function in SAC signalling and kinetochore-microtubule attachment^56,57^. To further examine the role of CDK4/6 in maintaining the SAC, we assessed the phosphorylation status of pMELT and BubR1 via pMELT and BubR1-specific anbibodies^58^. CDK4/6 inhibition substantially reduced pMELT levels, through the reduction was less pronounced compared to alisertib treatment (Fig. 3d, left). BubR1 bands showed a mobility shift indicative of dephosphorylation (Fig. 3d, left). Immunostaining with the pMELT-specific antibody^58^ corroborated these findings, showing reduced pMELT signals at kinetochores following treatment with CDK4/6 inhibitors (Fig. 3d, right). Notably, despite this reduction, pMELT signals remained detectable at kinetochores even after 7 hours of CDK4/6 inhibition (Fig. 3e), indicating that CDK4/6 inhibition does not entirely abolish SAC activity, leaving residual signallling at unattached kinetochores. Together, these results suggest that CDK4/6 activity is crucial for sustaining robust SAC signalling and checkpoint integrity for prolonged mitotic arrest.

### CDK4 associates with chromosomes during early mitosis and SAC-dependent mitotic arrest

Our findings reveal a novel mitotic role for CDK4/6 in modulating the SAC. To elucidate the underlying mechanism, we investigated the behaviour of CDK4 and CDK6 during mitosis in HCT116 cells. Although CDK4/6 expression levels are thought to be constant throughout the cell cycle^59^, immunostaining analyses demonstrated dynamic localization patterns of these kinases during mitosis (Extended Data Fig. 5). Both CDK4 and CDK6 predominantly localised to the nucleus during interphase and became dispersed as cells entered mitosis (Extended Data Fig. 5a, b). Notably, CDK4 maintained a weak association with chromosomes during early mitosis (Extended Data Fig. 5a), a localisation that was more pronounced in cells arrested in prometaphase through SAC activation via nocodazole treatment (Extended Data Fig. 5c, d). This chromosome association suggests a potential direct role for CDK4 in mitotic regulation.

Among the known CDK4/6 cyclin partners, Cyclin D1 (CycD1) exhibited a localisation pattern similar to that of CDK4 (Extended Data Fig. 5e). It predominantly accumulated in the interphase nucleus but also showed a weak association with chromosomes during early mitosis (Extended Data Fig. 5e, Prophase). However, compared to CDK4/6, CycD1 levels varied significantly between cells, likely reflecting differences in mitogenic signals and cellular stress (Extended Data Fig. 5e, compare "High" and "Low"). This variability suggests that while CycD1 likely partners with CDK4 on mitotic chromosomes, its expression and, consequently, mitotic CDK4 activity, may be dynamically modulated by both external and internal cues.

### CDK4/6 regulate phosphorylation of the mitotic apparatus, including SAC components

To elucidate the molecular mechanisms by which CDK4/6 regulate the SAC, we conducted phosphoproteomic analyses of mitotic HCT116 cells treated with palbociclib (see Methods). Cells were first arrested in mitosis using nocodazole and subsequently treated with 1 µM or 5 µM palbociclib for 2 hours, an interval sufficient to induce BubR1 dissociation from kinetochores (Fig. 3c, Fig. 4a). MG132 was also added to prevent phosphoproteomic alterations associated with mitotic exit (Fig. 4a).

**Figure 4.**
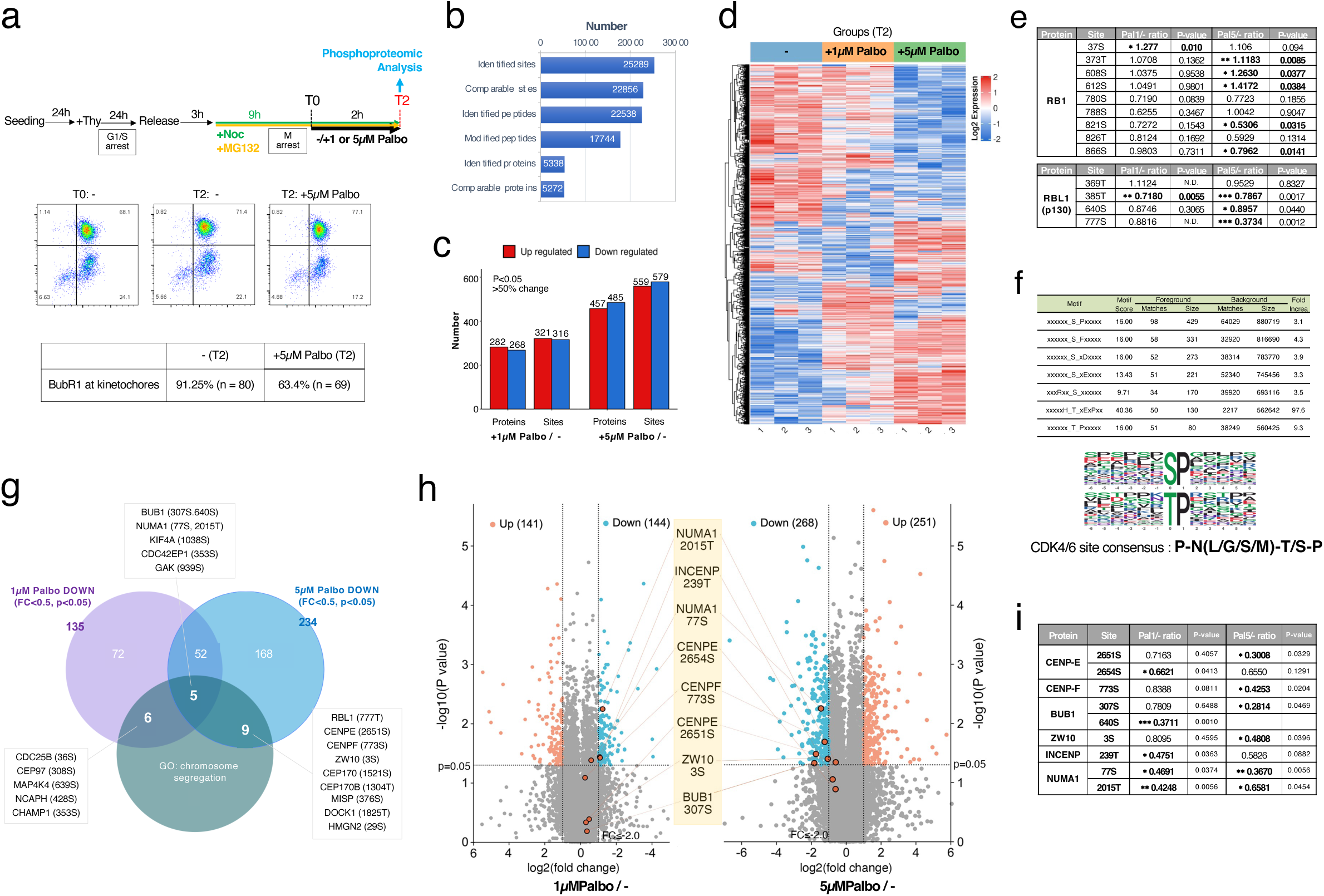
CDK4 regulates phosphorylation of mitotic apparatus, including SAC components. **a,** Schematic of the experimental workflow for phosphoproteomic analysis. HCT116 cells were synchronised at prometaphase by 9-hour nocodazole and MG132 treatment after thymidine release (T0). Cells were exposed to 1 µM or 5 µM Palbo or control (-) for 2 hours (T2) and harvested for phosphoproteomic analysis. BubR1 dissociation from kinetochores and cell cycle profiles were confirmed as indicators of CDK4/6 inhibition efficiency. **b,** Summary of identified phosphorylation sites, peptides, and proteins across all samples. Sites and proteins that were comparable between treatments are shown. **c,** Numbers of differentially phosphorylated sites (upregulated and downregulated) and proteins in 1 µM and 5 µM Palbo-treated cells compared to controls (fold change > 1.5 or < 0.667, p < 0.05). **d,** Heatmap of global phosphorylation profiles across all samples, showing changes induced by 1 µM and 5 µM Palbo treatment. **e,** Established CDK4/6 substrates, RB1 and RBL1, showed significant reduction in phosphorylation upon treatment with Palbo treatment. **f,** Motif enrichment analysis of differentially phosphorylated sites revealed significant enrichment of the pS/T-P motif, characteristic of CDK4/6 substrates. **g,** Venn diagram illustrating chromosome segregation-related proteins among significantly downregulated phosphoproteins upon 1 µM or/and 5 µM Palbo treatment (fold change < 0.5, p < 0.05). **h,** Volcano plots showing differentially phosphorylated proteins associated with chromosome segregation in 1 µM and 5 µM Palbo treatments compared to controls. Key SAC-related proteins are highlighted. **i,** Putative CDK4/6 phosphorylation sites identified on SAC-related proteins, CENP-E, CENP-F, BUB1, ZW10, INCENP, and NUMA1, with significant reduction in phosphorylation upon Palbo treatment.

In total, we identified 25,289 phosphorylation sites on 5,338 proteins, of which 22,856 sites on 5,272 proteins were quantifiable across conditions (Fig. 4b, Extended Data Table 1). Compared to control cells, 321 and 316 phosphorylation sites on 282 and 268 proteins, respectively, were significantly upregulated, while 559 and 579 sites on 457 and 485 proteins, respectively, were significantly downregulated (fold change > 1.5 or < 0.667, p < 0.05) following 1 µM or 5 µM palbociclib treatment (Fig. 4c, d). Notably, downregulated phosphorylation sites included those on known CDK4/6 targets such as RB1 and RBL1 (p107) (Fig. 4e), supporting the robustness of this approach to identify CDK4/6 substrates potentially involved in SAC regulation. Motif enrichment analysis for the phosphorylation sites significantly downregulated in Palbo-treated cells revealed a significant enrichment of the pS/T-P motives (Motif score: 16.00), including the P-x-pT/S-P motif, which is characteristic of CDK4/6 substrates (Fig. 4f). This strongly suggests that the observed phosphorylation changes are a direct consequence of CDK4/6 inhibition.

Next, we focused on phosphorylation sites with a reduction of over 50% (Log_2_FoldChange < −1) following treatment with 1 µM or 5 µM palbociclib and identified sites associated with chromosome segregation (Fig. 4g, Extended Data Table 2). From this analysis, we identified six proteins, CDC25B, CHAMP1, CEP97, MAP4K4, INCENP, and NCAPH, with significantly reduced phosphorylation in the 1 µM treatment condition, nine proteins, RBL1, CENPE, CENPF, ZW10, CEP170, CEP170B, MISP, DOCK1, and HMGN2, in the 5 µM condition, and five proteins, BUB1, NUMA1, KIF4A, CDC42EP1, and GAK, with reductions under both treatment conditions (Fig. 4g). Of these, known components and regulators of the SAC, including CENP-E, CENP-F, BUB1, ZW10, and INCENP, were identified as significantly downregulated in either or both conditions (Fig. 4h, i). These findings indicate that CDK4/6 regulate the phosphorylation status of proteins in SAC signalling and chromosome segregation during mitosis.

### CDK4-CycD1 phosphorylates SAC components including CENP-E

We assessed whether CDK4/6 directly phosphorylates these SAC components using *in vitro* kinase assays. Recombinant Cdk4-CycD1 complexes were incubated with purified fragments of the candidate proteins in the presence of ATP-γ-S. Gollowing alkylation, thiophosphates were detected using anti-thiophosphate antibodies^60^ (see Methods). The Cdk4-CycD1 complex efficiently phosphorylated its known substrates, RB (amino acids 380-928) and RBL1 (391-972) (Fig. 5a). Among the candidate mitotic substrates, CENP-E (2452-2701), CENP-F (2831-3114), and Bub1 (300-479) were phosphorylated by Cdk4-CycD1 (Fig. 5a-c).

**Figure 5.**
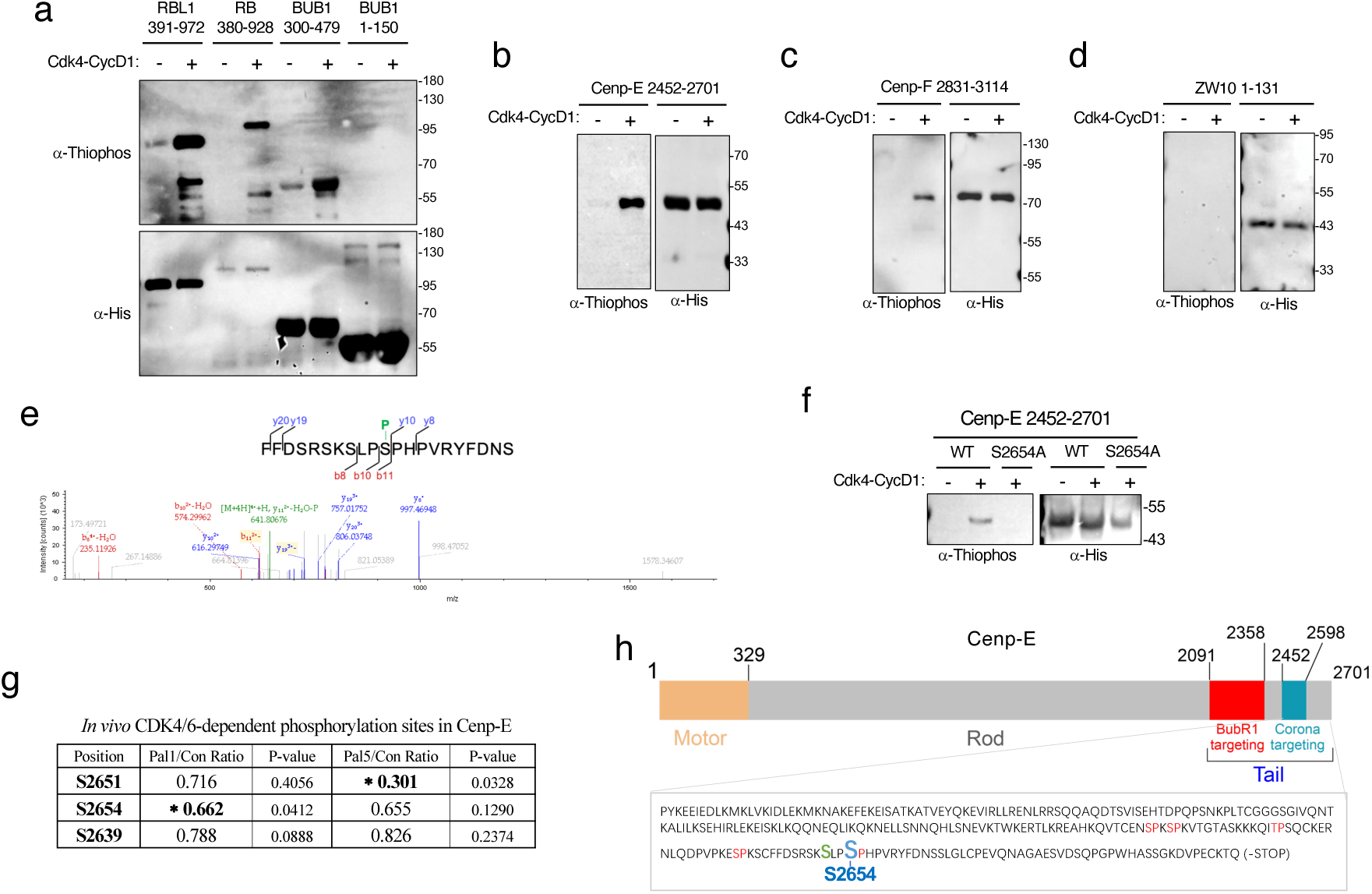
CDK4 directly phosphorylates critical SAC regulators, including CENP-E. **a,** *In vitro* kinase assays show that recombinant Cdk4–CycD1 complexes phosphorylate fragments of Bub1 (300–479), CENP-E (2452–2701), and CENP-F (2831–3114), but not Bub1 (1–150) or ZW10 (1–131). RBL1 (391–972) and RB (380–928) served as positive controls. Phosphorylation was detected using ATP-γS with thiophosphate ester immunoblotting. Substrate input was validated using anti-His tag antibodies. **b-d,** Cdk4 phosphorylation of individual SAC components: **b,** CENP-E 2452–2701 showed robust phosphorylation. **c,** CENP-F 2831–3114 was similarly phosphorylated. **d,** ZW10 1–131 was not phosphorylated. **e,** Mass spectrometry analysis identifies serine 2654 (S2654) as the CDK4 phosphorylation site within the CENP-E 2452–2701 fragment. The annotated spectrum shows y- and b-ions confirming phosphorylation at S2654. **f,** *In vitro* phosphorylation assays with wild-type (WT) and S2654A mutant CENP-E fragments show abolished phosphorylation in the S2654A mutant, confirming S2654 as the CDK4-specific site. **g,** Phosphoproteomic analysis of Palbo-treated mitotic cells identifies S2654 among phosphorylation sites significantly reduced upon CDK4 inhibition (*: p < 0.05). **h,** Schematic representation of the CENP-E domain structure, showing motor, rod, and tail regions. The tail, enriched with S/T-P motifs, contains S2654, a CDK4 phosphorylation site crucial for BubR1 recruitment to kinetochores and SAC maintenance.

Mass spectrometry analysis following *in vitro* kinase assays with normal ATP, combined with mutational analysis, identified serine 2654 (S2654) within the CENP-E fragment as a CDK4 phosphorylation site (Fig. 5e). Phosphoproteomic analysis detected a consistent reduction of phosphorylation at S2654 and its neighbouring S2651 residue after treatment with 1µM and 5 µM palbocilib (Fig. 4h, i, Fig. 5g). The C-terminal domain of CENP-E, which includes S2654, is highly enriched in S/T-P motifs and is critical for its interaction with BubR1 and the recruitment of BubR1 to kinetochores, key processes for SAC maintenance^7,8^ (Fig. 5h). These findings strongly suggest that CDK4/6 regulates SAC activity by directly phosphorylating key SAC components such as CENP-E.

Altogether, our findings establish CDK4/6 as a crucial regulator of SAC function and chromosome segregation, providing mechanistic insights into its role in maintaining mitotic fidelity through phosphorylation of key substrates.

## Discussion

Our study uncovers an unexpected layer of complexity in the regulatory network linking mitogenic cues to mitotic fidelity. While CDK4/6 are traditionally viewed as central drivers of the G1–S transition, here we identify a critical function for CDK4/6 in sustaining the SAC and ensuring proper chromosome segregation. By inhibiting CDK4/6 activity after S phase initiation, we observed that cells prematurely exited mitosis despite unattached kinetochores, resulting in chromosome missegregation and the accumulation of cells with abnormal nuclear morphology (Fig. 1). Thus, contrary to the prevailing model that places CDK4/6 function upstream and limited to G1, we demonstrate that CDK4/6 activity is critical for maintaining the fidelity of mitotic progression.

One key insight from our work is that CDK4/6 inhibition attenuates but does not completely abolish SAC signalling. The partial reduction in phosphorylation of SAC components and the residual, albeit weakened, kinetochore-associated signalling (Fig. 3d) highlight a scenario in which CDK4/6 help to prolong and reinforce SAC-mediated mitotic arrest rather than directly initiate it. This perspective is reminiscent of observations from other cell cycle transitions, where small perturbations in kinase or phosphatase activities do not eliminate signal transduction but modulate its strength and duration^61,62^. By positioning CDK4/6 as modulators of the SAC, our findings bridge the gap between early proliferative signals and late mitotic surveillance mechanisms, ensuring that cells not only commit to division but also safeguard genomic integrity during that process.

Our phosphoproteomic and *in vitro* phophorylation analysis identified a collection of CDK4/6-dependent phosphorylation events on key mitotic regulators, including CENP-E (Fig. 4, 5). The identification of a CDK4 phosphorylation site in the CENP-E tail domain, which is crucial for BubR1 recruitment and SAC maintenance^7,8^, provides a mechanistic anchor for understanding how CDK4/6 influence SAC activity. Although the SAC can be established through canonical pathways involving Mps1 and Aurora B kinases^20,21^. our data indicate that CDK4/6 fine-tune these checkpoint signals to ensure accurate chromosome alignment and tension. This nuanced control might be particularly important in cells experiencing fluctuating external signals or stress conditions, allowing them to adjust the threshold for mitotic arrest and chromosome segregation fidelity in response to their proliferative state.

Our findings also have implications for cancer biology and therapy. Cancer cells often exhibit compromised SAC function and tolerance to chromosomal instability, fueling tumour evolution and resistance to chemotherapies^29–31^. Here, we show that CDK4/6 inhibition can further weaken SAC arrest, promoting mitotic slippage and aneuploid. While this outcome may seem detrimental, it also unveils a potential therapeutic opportunity. Combining CDK4/6 inhibitors with SAC-targeting agents, such as Mps1 inhibitors, may synergistically push cells towards lethal chromosomal missegregation and apoptosis^30^. This combination strategy might enhance the efficacy of current anti-mitotic drugs, which often fail due to the cell’s ability to slip out of prolonged mitotic arrest^22,23^. In this way, CDK4/6 inhibition could sensitize cancer cells to therapies that exploit their mitotic vulnerabilities, even beyond the well-established G1 arrest currently leveraged in the clinic^4,5^.

Future studies will be needed to dissect the full range of CDK4/6 substrates at mitosis and to understand how changes in mitogenic signalling, metabolic states, or stress responses influence these newly appreciated roles in mitotic control.

## Extended Data

**Extended Data Figure 1 Effects of CDK4/6 inhibition on cell cycle distribution and Rb phosphorylation.**

Asynchronous HCT116, MCF7, and RPE-1 hTERT cells were treated with CDK4/6 inhibitors (CDK4/6-in) or vehicle control for up to 48 hours. DNA content was analysed by flow cytometry, and phosphorylated Rb at serine 780 (pRb) levels were examined by Western blot. **a**, Schematic representation of the experimental time-course setup. **b**, Flow cytometry analysis of cell cycle profiles (G1, S, G2/M) in HCT116 cells treated with increasing concentrations of palbociclib (Palbo). **c**, Representative immunoblot showing pRb, total Rb, and β-tubulin (β-Tub) levels in HCT116 cells treated with Palbo for 24 hours (top). Quantification of pRb band intensities normalized to total Rb is plotted, with IC₅₀ and C_max_ values indicated (bottom). **d, e**, HCT116 cells were treated with increasing concentrations of Palbo, abemaciclib (Abema), ribociclib (Ribo), or the MEK inhibitor PD0325901 (MEKi) for 24 hours. **d**, Flow cytometry analysis of cell cycle profiles. **e**, Western blot analysis of pRb, total Rb, and β-tubulin levels. Quantified pRb levels normalized to total Rb are shown in bar graphs. **f**, MCF7 cells treated with Palbo were analysed by flow cytometry for cell cycle profiles at 24 hours. **g, h**, RPE-1 hTERT cells treated with Palbo were analysed. **g**, Flow cytometry analysis of DNA content. **h**, Western blot analysis of pRb, total Rb, and β-tubulin. Quantified pRb levels are plotted with IC₅₀ and Cₘₐₓ values estimated.

**Extended Data Figure 2 CDK4/6 inhibitors induce premature mitotic exit in nocodazole-arrested cells.**

**a,** Experimental scheme: HCT116 cells were synchronised at the G1/S boundary, released into nocodazole-containing medium for 9 hours to arrest in prometaphase, and treated with CDK4/6 inhibitors or MEK inhibitor (MEKi). Samples were collected at the indicated time points for analysis. **b,** Relative mitotic populations (4N DNA content, pH3-positive) following treatment with increasing concentrations of palbociclib, abemaciclib, ribociclib, or MEKi were quantified by flow cytometry and plotted as arbitrary units, with the mitotic population at T0 set as 100. CDK4/6 inhibitors induced a dose-dependent reduction in mitotic cells, with abemaciclib being the most potent. **c,** Representative Western blot images showing phosphorylated Rb (pRb), total Rb, Cyclin B1 (CycB1), Cyclin D1 (CycD1), and β-tubulin (loading control) levels after treatment with CDK4/6 inhibitors or MEKi. **d,** Quantification of pRb and CycB1 band intensities from Western blot analyses. Relative levels at each time point were normalised to loading controls and plotted as arbitrary units. **e,** Comparative kinetics of mitotic exit induced by CDK4/6 inhibitors and MEKi based on mitotic populations at T3 (3 hours after addition). Abemaciclib induced the fastest mitotic exit, followed by palbociclib and ribociclib, while MEKi showed no significant effect.

**Extended Data Figure 3 CDK4/6 inhibition induces premature mitotic exit in diverse cell lines.**

**a, b,** Mitotic slippage following CDK4/6 inhibition was assessed in HCT116, HeLa, MCF7, and RPE-1 hTERT cells. Cells were treated with nocodazole for 24 hours to induce mitotic arrest (4N DNA content with pH3 signals). Palbo was added at the indicated concentrations, and samples were collected at specified time points for flow cytometry analysis. **a,** HCT116 and HeLa cells were treated with Palbo at 0, 0.5, 1, 2.5, 5, or 10 μM, and harvested at 0, 3, and 7 hours post-treatment. Line graphs show the proportion of mitotic cells (4N DNA content with positive pH3 signals). **b,** MCF7 and RPE-1 hTERT cells were treated with Palbo at 0, 1, 3, or 5 μM, and harvested at 0, 3, 5, 8, and 12 hours post-treatment. Similar to HCT116 cells, Palbo induced accelerated mitotic slippage in all cell lines, although the sensitivity to Palbo varied among the lines.

**Extended Data Figure 4 CDK4/6 inhibition-induced premature mitotic exit is independent of APC/CCDH1 activity and Aurora B kinase.**

**a,** Premature mitotic exit upon CDK4/6 inhibition is not mediated by APC/C^CDH1^. HCT116 cells expressing OsTIR1 and mAID-mCherry-tagged CDH1 were synchronised in prometaphase with nocodazole treatment. Depletion of CDH1 was induced by the addition of 5-Ph-IAA (500 µM), reducing CDH1 levels to below 20% after 3 hours (T0). Palbo was added at T0, and samples were collected at the indicated time points. Flow cytometry and Western blot analysis showed that mitotic exit and CycB1 degradation occurred with comparable kinetics in CDH1-depleted and control cells, indicating that CDH1 is not required for mitotic exit induced by CDK4/6 inhibition. **b,** Aurora B inhibition by alisertib triggers mitotic exit with loss of pH3 signals, while CDK4/6 inhibition does not affect pH3 levels in mitotic cells. HCT116 cells were treated with Palbo or alisertib after nocodazole-induced mitotic arrest. Immunofluorescence staining revealed that alisertib reduced pH3 signals despite the presence of condensed mitotic chromosomes, whereas Palbo-treated cells retained pH3 signals. Scale bar: 10 µm. Quantification of pH3-positive and -negative mitotic cells is shown as a bar graph.

**Extended Data Figure 5 CDK4 associates with chromosomes during early mitosis and SAC arrest.**

**a, b,** Subcellular localisation of CDK4 and CDK6 was examined in asynchronous HCT116 cells using immunofluorescence staining. CDK4 and CDK6 predominantly localised to the nucleus during interphase and were dispersed as cells progressed through mitosis. CDK4 exhibited a weak association with chromosomes during early mitosis (prophase to prometaphase). Chromosome morphology and pH3 staining were used to determine cell cycle phases. Scale bar: 10 µm. **c, d,** Localisation of CDK4 during SAC arrest. HCT116 cells were synchronised at G1/S by a thymidine block, released into nocodazole-containing medium, and harvested after 9 hours to arrest in prometaphase. Immunofluorescence staining revealed pronounced CDK4 association with chromosomes during SAC-dependent mitotic arrest. Aurora kinase B (AurB) served as an inner kinetochore marker. Scale bar: 10 µm. **e,** Cyclin D1 (CycD1) showed nuclear localisation in interphase and weak association with chromosomes during early mitosis (prophase to prometaphase). CycD1 expression levels varied between cells, reflecting fluctuations in mitogenic signals. pH3 staining indicated mitotic cells. Scale bar: 10 µm.

**Extended Data Table 1 Identified phosphorylation sites.**

A comprehensive list of 25,289 phosphorylation sites identified across 5,338 proteins in HCT116 cells. Of these, 22,856 sites across 5,272 proteins were quantifiable in the conditions tested (control, 1 µM palbociclib, and 5 µM palbociclib; Fig. 4b). Each entry includes the protein accession number, phosphorylation position, amino acid modification, protein description, gene name, modified peptide sequence, localisation probability, and quantitative intensity values for each condition and replicate. Subcellular localisation, KEGG pathway annotations, associated biological processes, and functional domains are also provided.

**Extended Data Table 2 Differentially phosphorylated sites.**

Phosphorylation sites exhibiting a reduction of over 33% in response to 1 µM or 5 µM palbociclib treatment, as well as those associated with chromosome segregation (Fig. 4g). Entries include protein accession number, position, regulated type (up or downregulated), phosphorylation site intensity values for control and palbociclib-treated samples, and fold changes with statistical significance (P values). The table also lists protein descriptions, gene names, modified peptide sequences, localisation probabilities, and functional annotations such as subcellular localisation, KEGG pathway involvement, biological processes, and structural domains.

## Methods

### Cell culture

Human HCT116 colon carcinoma cellswere maintained in Dulbecco’s Modified Eagle Medium (DMEM; Gibco, 11965-092) supplemented with 10% fetal bovine serum (FBS; LONSERA, S711-001) and Penicillin-Streptomycin (Gibco, 15140-122) at a concentration recommended by the manufacturer. Cultures were incubated at 37°C in a humidified atmosphere of 5% CO_2_ and were routinely passaged at 80–95% confluence to prevent overgrowth. The culture medium was replaced every 2–3 days. In addition, MCF7 breast carcinoma cells, and HeLa cervical carcinoma cells, and RPE1 hTERT-immortalised retinal pigment epithelial cellswere also cultured with modifications specific to each cell line.

### Cell Synchronization

G1/S synchronisation was achieved using modified thymidine block protocols based on Schvartzman et al.^63^ and Bostock et al.^64^ For a single thymidine block, cells were seeded at approximately 20–30% density per well and allowed to adhere for 24 hours until reaching ∼50% confluence. Culture medium was then replaced with DMEM containing 2 mM thymidine (Sigma-Aldrich, T1895-5G) for 24 hours. After this incubation, cells were washed three times with pre-warmed PBS and released into regular growth medium.

For the double thymidine block, cells were first incubated with 2 mM thymidine at ∼40% confluence for 18 hours, washed twice with PBS, and then released into fresh medium at 37°C for 9 hours. A second 2 mM thymidine treatment was applied for an additional 15 hours, followed by two PBS washes and release into fresh medium. Both single and double thymidine block protocols typically resulted in >90% synchronization at the G1/S boundary. Mitotic arrest was induced by releasing G1/S-arrested cells into standard growth medium for 3 hours to complete S phase and enter G2/M. The medium was then replaced with DMEM containing 100 ng/mL nocodazole (MCE, HY-13520) for 9–12 hours. Under these conditions, approximately 70–80% of cells were arrested in prometaphase. Cells were collected at defined time points to match the experimental requirements of subsequent assays.

### Pharmacological treatments

Following synchronisation, cells were treated with various inhibitors. To prevent proteasomal degradation, MG132 (Selleck, 1211877-36-9; MCE, HY-13259C) was used. For kinase inhibition, we employed CDK4/6 inhibitors—palbociclib (MCE, HY-50767), abemaciclib (MCE, HY-16297A), and ribociclib (MCE, HY-15777)—as well as the CDK1-specific inhibitor RO3306 (Selleck, 872573-93-8; MCE, HY-12529), the Aurora A kinase inhibitor alisertib (MCE, HY-10971) and the MEK inhibitor PD0325901 (Selleck S1036).). Concentrations and treatment durations for each inhibitor are detailed in the respective figure legends.

### Flow Cytometry

Cells were collected on ice to minimise metabolic activity. After washing twice with ice-cold PBS, cells were trypsinised, pelleted (1, 500 rpm, 5 min, 4°C), and washed with 1% BSA/PBS (wash buffer). For fixation, cell pellets were resuspended in 4% paraformaldehyde at room temperature for 15 min, then washed and stored in wash buffer at 4°C for up to three days.

For staining, fixed cells were permeabilized with 5% Triton X-100/PBS (permeabiliasation buffer) for 15 min at room temperature in the dark. For pH3 and Cyclin B detection, phospho histone H3 Ser10 antibodies (Invitrogen, PA5-17869, 1:300) and Cyclin B1 antibodies (Santa-Cruz, sc-245, 1:100) (diluted in permeabilization buffer) were applied for 30 min at room temperature, followed by a wash and a 30-min incubation with secondary antibodies including Goat Anti-Mouse secondary antibody, Alexa Fluor® 488 (Abcam, ab150113, 1:1000), Goat Anti-Rabbit secondary antibody, Alexa Fluor® 568 (Abcam, ab175471, 1:1000), Goat anti-Guinea Pig secondary antibody, Alexa Fluor™ 647 (Invitrogen, A-21450, 1:1000). After a final wash, DNA was stained using Hoechst 33342 (10 μg/mL, APExBio, A3472) or Propidium Iodide (PI, ab139418, Abcam) following the manufacturer’s instructions.

All samples were filtered through a 74 μm mesh before analysis. Negative controls (no primary antibody) and positive controls (metaphase-arrested cells) were included. Flow cytometry was performed using a BD LSRFortessa cytometer, collecting FSC, SSC, DNA, and fluorescence parameters. Data were analysed with FlowJo software using standard gating strategies (FSC-A/SSC-A, FSC-A/SSC-W, DNA-A/DNA-W) to identify single live cells and the desired cell populations. DNA content and target protein levels were quantified and further analysed using GraphPad.

### Western Blot

Cells were lysed directly in 2× Laemmli sample buffer or in RIPA buffer supplemented with protease and phosphatase inhibitors. Lysates were clarified by centrifugation, normalised by protein concentration, denatured at 95°C, and resolved by SDS-PAGE. Proteins were transferred to PVDF membranes, blocked in 5% milk or BSA in TBST, and probed with primary antibodies at 4°C overnight, followed by HRP-conjugated secondary antibodies for 1 hour at room temperature. After washing, membranes were developed with chemiluminescent reagents and imaged. Antibodies used for Western blotting included phospho-Rb (CST #8180), Rb (CST #9309), β-Tubulin (ABclonal A12289), BubR1 (Abclonal A1775), Cdc27 (Abcam Ab10538), Cyclin B1 (Santa Cruz sc-245), Cyclin D1 (Santa Cruz sc-8396), GAPDH (Abcam 9484), KNL1 (Abcam ab70537), CDH1/FZR1 (Santa Cruz sc-166714), Mps1/TTK (Abclonal A2500), CDK4 (Santa Cruz sc-23896), CDK6 (Santa Cruz sc-7961), phospho-KNL1 MELT (CST #40758), thiophosphate ester (Abcam ab92570), and HRP-conjugated His-tag antibodies (Abclonal AE104). Species-specific HRP-conjugated secondary antibodies (Thermo Fisher) were used at 1:10,000.

### Immunofluorescence

Cells grown on coverslips were fixed with either cold methanol or 4% paraformaldehyde, permeabilised with Triton X-100, and blocked in 5% goat serum (Beyotime C0265). Primary antibodies were incubated overnight at 4°C, followed by fluorescently conjugated secondary antibodies for 1 hour at room temperature. DNA was stained with Hoechst 33342 (APExBIO A3472). Coverslips were mounted with antifade mounting medium and imaged by Nikon Ti2 wide-field microscopy or C2 confocal fluorescence microscopy.

Antibodies used for immunofluorescence included Lamin A/C (Abcam ab238303), β-Tubulin (ABclonal A12289), BubR1 (Abclonal A1775), CENP-C (MBL PD030), phospho-Histone H3 Ser10 (ABclonal AP0002), phospho-KNL1 MELT (CST #40758), CDK4 (Santa Cruz sc-23896), and CDK6 (Santa Cruz sc-7961), and Cyclin D1 (Santa Cruz sc-8396). Appropriate Alexa Fluor-conjugated secondary antibodies (Abcam, Invitrogen) were used at 1:1000.

Images were processed using ImageJ, and co-localization or intensity measurements were quantified with ImageJ plugins and graphed with GraphPad Prism.

Colocalisation analysis was performed using the Colocalization Finder plugin of ImageJ software which is based on a practical guide reported by Dunn et al^65^.

### Phosphoproteomic Analysis of Mitotic HCT116 Cells Following CDK4/6 Inhibition

Mitotically arrested HCT116 cells were prepared as described above. Briefly, cells were synchronized at the G1/S boundary, released into S phase, and arrested in mitosis with nocodazole. MG132 (MCE, HY-13259C) was added to prevent mitotic exit, and 1 µM or 5 µM palbociclib was introduced for 2 hours to inhibit CDK4/6 activity. Control samples were treated with vehicle only. For each condition, three biological replicates were collected. After treatments, cells were harvested on ice, washed with PBS, and pelleted by centrifugation. Pellets were snap-frozen in liquid nitrogen and stored at −80°C until analysis by Jingjie PTM Biotech (Hangzhou, China). Protein extraction, tryptic digestion, and phosphopeptide enrichment (e.g., TiO_2 or IMAC) were conducted following standard protocols. Phosphopeptides were analyzed by LC-MS/MS on a high-resolution mass spectrometer, and data were processed using software such as MaxQuant or Proteome Discoverer against a human proteome database. Quantitative comparisons identified phosphosites significantly altered by palbociclib treatment (p < 0.05). Motif enrichment analysis was performed using MoMo^66^ to detect enrichment of characteristic phosphorylation motifs (e.g., pS/T-P) in phosophorylation sites that showed significant reduction (fold change < 0.5) in Palbo-treated cells compared to control, and identified substrates were mapped to key mitotic regulators, including SAC components. Results were interpreted in the context of CDK4/6-dependent mitotic phosphorylation events.

### *In vitro* CDK4 Kinase Assay

Candidate SAC substrate fragments, including Bub1 1–150, 300–479, ZW10 1–131, CENP-E 2452–2701, and CENP-F 2831–3114 amino-acid fragments, were amplified from HCT116 cDNA and cloned into bacterial expression vectors, each containing N- or C-terminal His and either GST or MBP tags. Phosphorylation site mutations (e.g., CENP-E S2691A) were introduced in CENP-E by site-directed mutagenesis. Constructs were verified by DNA sequencing and expressed *in E. coli* BL21(DE3) cells. Cultures were induced with 0.1 mM IPTG at OD_600_ 0.6–1.0 and grown at 16°C for 16–24 hours. Cells were lysed in a buffer containing 20 mM Tris-HCl (pH 7.5), 0.5 M NaCl, and 5 mM imidazole, supplemented with protease inhibitors. Lysates were clarified and bound to Ni-NTA resin. After washing with increasing imidazole concentrations, proteins were eluted with 0.5–1.0 M imidazole, concentrated, and exchanged into kinase buffer (50 mM Tris-HCl, pH 7.5, 150 mM NaCl) supplemented with 20% glycerol before storage at −80°C.

Purified substrates were incubated with an active recombinant human CDK4–Cyclin D1 complex (Abcam, ab55695) in the kinase buffer (25 mM Tris-HCl, pH 7.5; 10 mM MgCl_2_; 0.3 mM DTT) at 30°C for 2 hours. For standard phosphorylation, 1 mM ATP was used. Thiophosphorylation labeling was performed following Allen et al., 2007^60^. 1m M ATP-γ-S (Aladdin Scientific, A274887) replaced ATP. After the kinase reaction, proteins were alkylated with 2.5 mM p-nitrobenzyl mesylate (PNBM, Selleck, E1248) in 5% DMSO for 1– 2 hours at room temperature to lock in the thiophosphate ester modification.

Following alkylation, samples were mixed with 5× Laemmli buffer and heated at 95°C for 10 minutes. Proteins were resolved by SDS-PAGE and transferred to PVDF membranes. Thiophosphorylated substrates were detected by Western blot using a thiophosphate ester-specific antibody (Abcam, ab92570) at 1:5,000 dilution in 5% milk/TBST. Membranes were probed with HRP-conjugated secondary antibodies and visualised by chemiluminescence.

### Identification of CDK4-mediated *in vitro* Phosphorylation Sites

*In vitro* kinase assays were performed as described above, using purified substrate proteins, CENP-E 2452-2701, CENP-F 2831-3114 and Bub1 300-479 amino-acid fragments, active CDK4–Cyclin D1 and ATP. After the reaction, samples were separated by SDS-PAGE, followed by Coomassie Brilliant Blue staining. The bands corresponding to the substrates were excised from the gel, and in-gel tryptic digestion was conducted as described previously^67^. The resulting peptides were extracted, dried, and reconstituted in 0.1% formic acid for mass spectrometry. Peptides were separated and analysed on an Easy-nLC 1200 system coupled to an Orbitrap Fusion (Thermo Scientific). About 0.5 µg of peptides were separated in an home-made column (75 µm x 15 cm) packed with C18 AQ (5 µm, 300Å, Michrom BioResources, Auburn, CA, USA) at a flow rate of 250 nL/min. Mobile phase A (0.1% formic acid) and mobile phase B (0.1% formic acid in 80% ACN) were used to establish a 60 min gradient comprised of 50 min of 6-34% B, 3 min of 34-38% B, 1 min of 38-90% B and 6 min of 90% B. Peptides were then ionized by electrospray at 2.1 kV. A full MS spectrum (350-1400 m/z range) was acquired at a resolution of 120,000 at m/z 200 and a maximum ion accumulation time of 50 ms. Dynamic exclusion was set to 30 s. Resolution for HCD MS/MS spectra was set to 15,000 at m/z 200. The AGC setting of MS and MS^2^ were set at 5E5 and 5E4, respectively. Isolation width of 1.6 m/z units and a maximum ion accumulation time of 50 ms were used for MS^2^. Single and unassigned charged ions were excluded from MS/MS. For HCD, normalized collision energy was set to 28%. The raw data were processed and searched with MaxQuant 1.6.5.0 with MS tolerance of 4.5 ppm, and MS/MS tolerance of 20 ppm. The UniProt mouse protein database (release 2016_07, 49863 sequences) and database for proteomics contaminants from MaxQuant were used for database searches. Reversed database searches were used to evaluate false discovery rate (FDR) of peptide and protein identifications. Two missed cleavage sites of trypsin were allowed. Oxidation (M), Acetyl (Protein N-term) and Deamidation (NQ) were set as variable modifications. The FDR of both peptide identification and protein identification is set to be 1%^68^. The option of “Second peptides” and “Match between runs” was enabled^75,76^. As a result, phosphorylation on serine 2654 of CENP-E and on serine 2911 of CENP-F were detected. No phosphorylation site was detected within the Bub1 300-479 amino-acid fragment.

## Acknowledgements

The authors thank Drs. Geert Kops, Randy Poon, Mingwei Min, Junseog Kang, Wei Qi, Cang Yong, and Zhenge Luo for advice and sharing reagents, and ShanghaiTech Cell Molecular Bilogy Core, Multiomics Core, and Imaging Core for their services and technical advice. The authors also thank all Kimata’s lab members for discussion. This work is supported by start-up funding from ShanghaiTech University (2018F0202-000-06), National Scientific Foundation of China (NSFC) Research Fund for International Scientist (RFIS, 32150710520) and Mianshang Project (32170746) to YK, and Science and Technology Comision of Shanghai Municipality (STCSM) Foreign Experts (21WZ2502400, 22WZ2503200, 20WZ2504700) to YK and ML

## References

1 Rubin, S. M., Sage, J. & Skotheim, J. M. Integrating Old and New Paradigms of G1/S Control. Mol Cell 80, 183–192 (2020). 10.1016/j.molcel.2020.08.020

2 Limas, J. C. & Cook, J. G. Preparation for DNA replication: the key to a successful S phase. FEBS Lett 593, 2853–2867 (2019). 10.1002/1873-3468.13619

3 Bertoli, C., Skotheim, J. M. & de Bruin, R. A. Control of cell cycle transcription during G1 and S phases. Nat Rev Mol Cell Biol 14, 518–528 (2013). 10.1038/nrm3629

4 Piezzo, M. et al. Targeting Cell Cycle in Breast Cancer: CDK4/6 Inhibitors. Int J Mol Sci 21 (2020). 10.3390/ijms21186479

5 Goel, S., DeCristo, M. J., McAllister, S. S. & Zhao, J. J. CDK4/6 Inhibition in Cancer: Beyond Cell Cycle Arrest. Trends Cell Biol 28, 911–925 (2018). 10.1016/j.tcb.2018.07.002

6 Musacchio, A. The Molecular Biology of Spindle Assembly Checkpoint Signaling Dynamics. Curr Biol 25, R1002–1018 (2015). 10.1016/j.cub.2015.08.051

7 Weaver, B. A. et al. Centromere-associated protein-E is essential for the mammalian mitotic checkpoint to prevent aneuploidy due to single chromosome loss. J Cell Biol 162, 551–563 (2003). 10.1083/jcb.200303167

8 Mao, Y., Abrieu, A. & Cleveland, D. W. Activating and silencing the mitotic checkpoint through CENP-E-dependent activation/inactivation of BubR1. Cell 114, 87–98 (2003). 10.1016/s0092-8674(03)00475-6

9 Matsushime, H. et al. Identification and properties of an atypical catalytic subunit (p34PSK-J3/cdk4) for mammalian D type G1 cyclins. Cell 71, 323–334 (1992). 10.1016/0092-8674(92)90360-o

10 Sherr, C. J. & Roberts, J. M. CDK inhibitors: positive and negative regulators of G1-phase progression. Genes Dev 13, 1501–1512 (1999). 10.1101/gad.13.12.1501

11 Wood, D. J. & Endicott, J. A. Structural insights into the functional diversity of the CDK-cyclin family. Open Biol 8 (2018). 10.1098/rsob.180112

12 McKenney, C. et al. CDK4/6 activity is required during G(2) arrest to prevent stress-induced endoreplication. Science 384, eadi2421 (2024). 10.1126/science.adi2421

13 Cornwell, J. A. et al. Loss of CDK4/6 activity in S/G2 phase leads to cell cycle reversal. Nature 619, 363–370 (2023). 10.1038/s41586-023-06274-3

14 Anders, L. et al. A systematic screen for CDK4/6 substrates links FOXM1 phosphorylation to senescence suppression in cancer cells. Cancer Cell 20, 620–634 (2011). 10.1016/j.ccr.2011.10.001

15 Wang, H. et al. The metabolic function of cyclin D3-CDK6 kinase in cancer cell survival. Nature 546, 426–430 (2017). 10.1038/nature22797

16 Goel, S. et al. CDK4/6 inhibition triggers anti-tumour immunity. Nature 548, 471–475 (2017). 10.1038/nature23465

17 Zhang, J. et al. Cyclin D-CDK4 kinase destabilizes PD-L1 via cullin 3-SPOP to control cancer immune surveillance. Nature 553, 91–95 (2018). 10.1038/nature25015

18 Foley, E. A. & Kapoor, T. M. Microtubule attachment and spindle assembly checkpoint signalling at the kinetochore. Nat Rev Mol Cell Biol 14, 25–37 (2013). 10.1038/nrm3494

19 Lara-Gonzalez, P., Westhorpe, F. G. & Taylor, S. S. The spindle assembly checkpoint. Curr Biol 22, R966–980 (2012). 10.1016/j.cub.2012.10.006

20 Lara-Gonzalez, P., Pines, J. & Desai, A. Spindle assembly checkpoint activation and silencing at kinetochores. Semin Cell Dev Biol 117, 86–98 (2021). 10.1016/j.semcdb.2021.06.009

21 McAinsh, A. D. & Kops, G. Principles and dynamics of spindle assembly checkpoint signalling. Nat Rev Mol Cell Biol 24, 543–559 (2023). 10.1038/s41580-023-00593-z

22 Rieder, C. L. & Maiato, H. Stuck in division or passing through: what happens when cells cannot satisfy the spindle assembly checkpoint. Dev Cell 7, 637–651 (2004). 10.1016/j.devcel.2004.09.002

23 Gascoigne, K. E. & Taylor, S. S. How do anti-mitotic drugs kill cancer cells? J Cell Sci 122, 2579–2585 (2009). 10.1242/jcs.039719

24 Mikwar, M., MacFarlane, A. J. & Marchetti, F. Mechanisms of oocyte aneuploidy associated with advanced maternal age. Mutat Res Rev Mutat Res 785, 108320 (2020). 10.1016/j.mrrev.2020.108320

25 Gerhold, A. R., Poupart, V., Labbé, J. C. & Maddox, P. S. Spindle assembly checkpoint strength is linked to cell fate in the Caenorhabditis elegans embryo. Mol Biol Cell 29, 1435–1448 (2018). 10.1091/mbc.E18-04-0215

26 Chen, S. Y., Cheng, P. W., Peng, H. F. & Wu, J. C. C. elegans spermatocyte divisions show a weak spindle checkpoint response. J Cell Sci 137 (2024). 10.1242/jcs.257675

27 Rebollo, E. & González, C. Visualizing the spindle checkpoint in Drosophila spermatocytes. EMBO Rep 1, 65–70 (2000). 10.1093/embo-reports/kvd011

28 Galli, M. & Morgan, D. O. Cell Size Determines the Strength of the Spindle Assembly Checkpoint during Embryonic Development. Dev Cell 36, 344–352 (2016). 10.1016/j.devcel.2016.01.003

29 Sarkar, S. et al. Mitotic checkpoint defects: en route to cancer and drug resistance. Chromosome Res 29, 131–144 (2021). 10.1007/s10577-020-09646-x

30 Sansregret, L., Vanhaesebroeck, B. & Swanton, C. Determinants and clinical implications of chromosomal instability in cancer. Nat Rev Clin Oncol 15, 139–150 (2018). 10.1038/nrclinonc.2017.198

31 Ben-David, U. & Amon, A. Context is everything: aneuploidy in cancer. Nat Rev Genet 21, 44–62 (2020). 10.1038/s41576-019-0171-x

32 Nabti, I., Marangos, P., Bormann, J., Kudo, N. R. & Carroll, J. Dual-mode regulation of the APC/C by CDK1 and MAPK controls meiosis I progression and fidelity. J Cell Biol 204, 891–900 (2014). 10.1083/jcb.201305049

33 Takenaka, K., Moriguchi, T. & Nishida, E. Activation of the protein kinase p38 in the spindle assembly checkpoint and mitotic arrest. Science 280, 599–602 (1998). 10.1126/science.280.5363.599

34 Wang, X. M., Zhai, Y. & Ferrell, J. E., Jr. A role for mitogen-activated protein kinase in the spindle assembly checkpoint in XTC cells. J Cell Biol 137, 433–443 (1997). 10.1083/jcb.137.2.433

35 Zhang, M., Kothari, P. & Lampson, M. A. Spindle assembly checkpoint acquisition at the mid-blastula transition. PLoS One 10, e0119285 (2015). 10.1371/journal.pone.0119285

36 Wu, X. et al. Distinct CDK6 complexes determine tumor cell response to CDK4/6 inhibitors and degraders. Nature Cancer 2, 429–443 (2021). 10.1038/s43018-021-00174-z

37 Hafner, M. et al. Multiomics Profiling Establishes the Polypharmacology of FDA-Approved CDK4/6 Inhibitors and the Potential for Differential Clinical Activity. Cell Chem Biol 26, 1067–1080.e1068 (2019). 10.1016/j.chembiol.2019.05.005

38 Liu, J. J. et al. Ras transformation results in an elevated level of cyclin D1 and acceleration of G1 progression in NIH 3T3 cells. Mol Cell Biol 15, 3654–3663 (1995). 10.1128/mcb.15.7.3654

39 Lavoie, J. N., L’Allemain, G., Brunet, A., Müller, R. & Pouysségur, J. Cyclin D1 expression is regulated positively by the p42/p44MAPK and negatively by the p38/HOGMAPK pathway. J Biol Chem 271, 20608–20616 (1996). 10.1074/jbc.271.34.20608

40 Scott, S. J., Suvarna, K. S. & D’Avino, P. P. Synchronization of human retinal pigment epithelial-1 cells in mitosis. Journal of Cell Science 133 (2020). 10.1242/jcs.247940

41 Bassermann, F., Eichner, R. & Pagano, M. The ubiquitin proteasome system - implications for cell cycle control and the targeted treatment of cancer. Biochim Biophys Acta 1843, 150–162 (2014). 10.1016/j.bbamcr.2013.02.028

42 Glotzer, M., Murray, A. W. & Kirschner, M. W. Cyclin is degraded by the ubiquitin pathway. Nature 349, 132–138 (1991). 10.1038/349132a0

43 Fujimitsu, K., Grimaldi, M. & Yamano, H. Cyclin-dependent kinase 1-dependent activation of APC/C ubiquitin ligase. Science 352, 1121–1124 (2016). 10.1126/science.aad3925

44 Morgan, D. O. Regulation of the APC and the exit from mitosis. Nature Cell Biology 1, E47–E53 (1999). 10.1038/10039

45 Fang, G., Yu, H. & Kirschner, M. W. Control of mitotic transitions by the anaphase-promoting complex. Philos Trans R Soc Lond B Biol Sci 354, 1583–1590 (1999). 10.1098/rstb.1999.0502

46 Zeng, X. & King, R. W. An APC/C inhibitor stabilizes cyclin B1 by prematurely terminating ubiquitination. Nat Chem Biol 8, 383–392 (2012). 10.1038/nchembio.801

47 Wan, L. et al. The APC/C E3 Ligase Complex Activator FZR1 Restricts BRAF Oncogenic Function. Cancer Discov 7, 424–441 (2017). 10.1158/2159-8290.Cd-16-0647

48 The, I. et al. Rb and FZR1/Cdh1 determine CDK4/6-cyclin D requirement in C. elegans and human cancer cells. Nat Commun 6, 5906 (2015). 10.1038/ncomms6906

49 Yesbolatova, A. et al. The auxin-inducible degron 2 technology provides sharp degradation control in yeast, mammalian cells, and mice. Nature Communications 11, 5701 (2020). 10.1038/s41467-020-19532-z

50 Courtheoux, T. et al. Aurora A kinase activity is required to maintain an active spindle assembly checkpoint during prometaphase. J Cell Sci 131 (2018). 10.1242/jcs.191353

51 Crosio, C. et al. Mitotic phosphorylation of histone H3: spatio-temporal regulation by mammalian Aurora kinases. Mol Cell Biol 22, 874–885 (2002). 10.1128/mcb.22.3.874-885.2002

52 London, N., Ceto, S., Ranish, J. A. & Biggins, S. Phosphoregulation of Spc105 by Mps1 and PP1 regulates Bub1 localization to kinetochores. Curr Biol 22, 900–906 (2012). 10.1016/j.cub.2012.03.052

53 Shepperd, Lindsey A. et al. Phosphodependent Recruitment of Bub1 and Bub3 to Spc7/KNL1 by Mph1 Kinase Maintains the Spindle Checkpoint. Current Biology 22, 891–899 (2012). 10.1016/j.cub.2012.03.051

54 Yamagishi, Y., Yang, C. H., Tanno, Y. & Watanabe, Y. MPS1/Mph1 phosphorylates the kinetochore protein KNL1/Spc7 to recruit SAC components. Nat Cell Biol 14, 746–752 (2012). 10.1038/ncb2515

55 Vleugel, M. et al. Arrayed BUB recruitment modules in the kinetochore scaffold KNL1 promote accurate chromosome segregation. J Cell Biol 203, 943–955 (2013). 10.1083/jcb.201307016

56 Huang, H. & Yen, T. J. BubR1 is an effector of multiple mitotic kinases that specifies kinetochore: Microtubule attachments and checkpoint. Cell Cycle 8, 1164–1167 (2009). 10.4161/cc.8.8.8151

57 Bloom, C. R. & North, B. J. Physiological relevance of post-translational regulation of the spindle assembly checkpoint protein BubR1. Cell Biosci 11, 76 (2021). 10.1186/s13578-021-00589-2

58 Vleugel, M. et al. Sequential Multisite Phospho-Regulation of KNL1-BUB3 Interfaces at Mitotic Kinetochores. Molecular Cell 57, 824–835 (2015). 10.1016/j.molcel.2014.12.036

59 Fassl, A., Geng, Y. & Sicinski, P. CDK4 and CDK6 kinases: From basic science to cancer therapy. Science 375, eabc1495 (2022). 10.1126/science.abc1495

60 Allen, J. J. et al. A semisynthetic epitope for kinase substrates. Nature Methods 4, 511–516 (2007). 10.1038/nmeth1048

61 Domingo-Sananes, M. R., Kapuy, O., Hunt, T. & Novak, B. Switches and latches: a biochemical tug-of-war between the kinases and phosphatases that control mitosis. Philos Trans R Soc Lond B Biol Sci 366, 3584–3594 (2011). 10.1098/rstb.2011.0087

62 Konagaya, Y., Rosenthal, D., Ratnayeke, N., Fan, Y. & Meyer, T. An intermediate Rb-E2F activity state safeguards proliferation commitment. Nature 631, 424–431 (2024). 10.1038/s41586-024-07554-2

63 Schvartzman, J. B., Krimer, D. B. & Van’t Hof, J. The effects of different thymidine concentrations on DNA replication in pea-root cells synchronized by a protracted 5-fluorodeoxyuridine treatment. Exp Cell Res 150, 379–389 (1984). 10.1016/0014-4827(84)90581-0

64 Bostock, C. J., Prescott, D. M. & Kirkpatrick, J. B. An evaluation of the double thymidine block for synchronizing mammalian cells at the G1-S border. Exp Cell Res 68, 163–168 (1971). 10.1016/0014-4827(71)90599-4

65 Dunn, K. W., Kamocka, M. M. & McDonald, J. H. A practical guide to evaluating colocalization in biological microscopy. Am J Physiol Cell Physiol 300, C723–742 (2011). 10.1152/ajpcell.00462.2010

66 Cheng, A., Grant, C. E., Noble, W. S. & Bailey, T. L. MoMo: discovery of statistically significant post-translational modification motifs. Bioinformatics 35, 2774–2782 (2019). 10.1093/bioinformatics/bty1058

67 Ren, Y. et al. The alterations of mouse plasma proteins during septic development. J Proteome Res 6, 2812–2821 (2007). 10.1021/pr070047k

68 Elias, J. E. & Gygi, S. P. Target-decoy search strategy for increased confidence in large-scale protein identifications by mass spectrometry. Nature Methods 4, 207–214 (2007). 10.1038/nmeth1019

69 Cox, J. & Mann, M. MaxQuant enables high peptide identification rates, individualized p.p.b.-range mass accuracies and proteome-wide protein quantification. Nat Biotechnol 26, 1367–1372 (2008). 10.1038/nbt.1511

70 Cox, J. et al. Andromeda: a peptide search engine integrated into the MaxQuant environment. J Proteome Res 10, 1794–1805 (2011). 10.1021/pr101065j

