## Supplementary figures for "Mitotic CDK4/6 activity sustains spindle checkpoint signalling to prevent mitotic slippage and genomic instability"

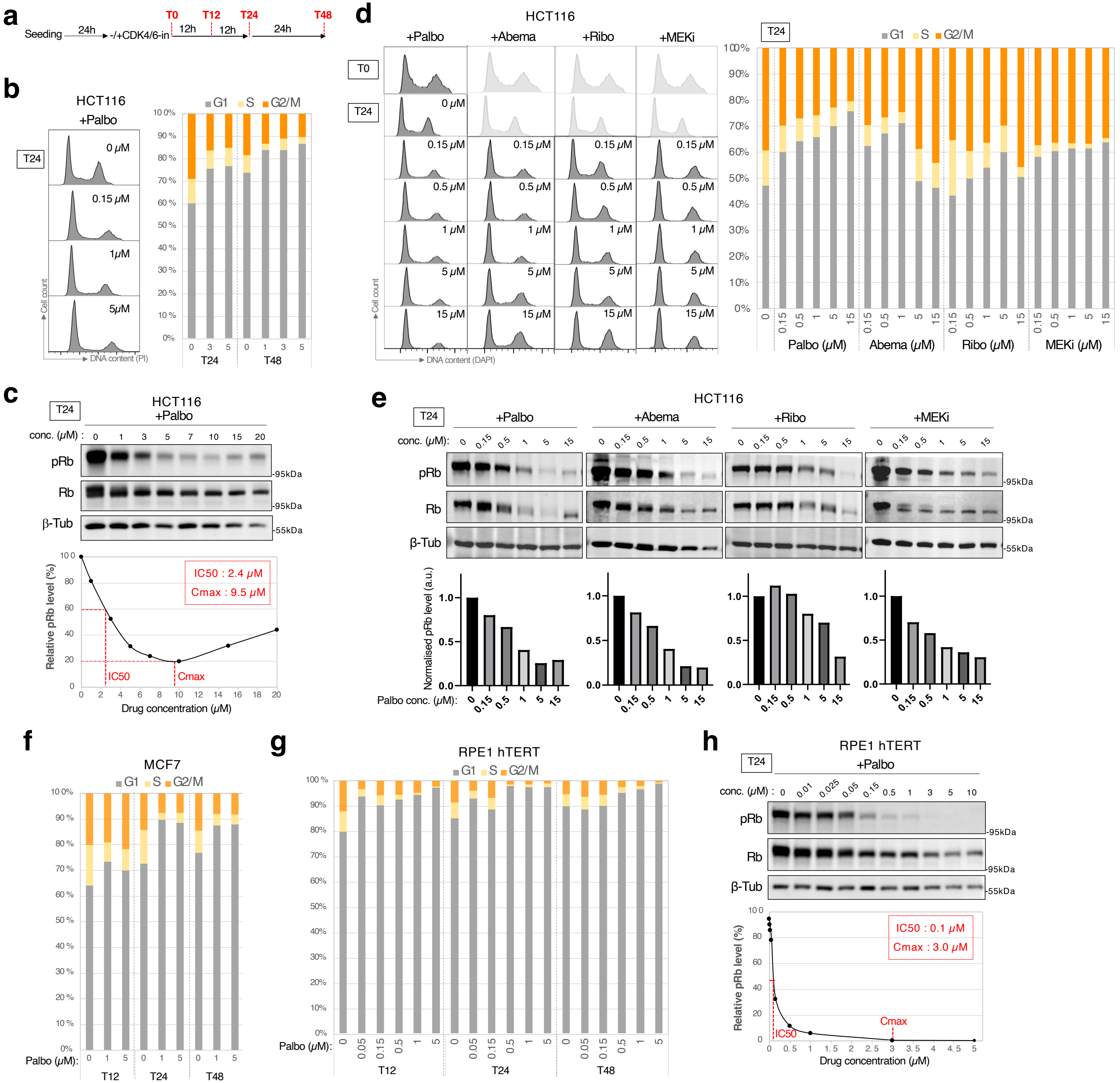


### Extended Data Figure 1 Effects of CDK4/6 inhibition on G1/S transition and Rb phosphorylation

**Extended Data**

**Extended Data Figure 1 Effects of CDK4/6 inhibition on cell cycle distribution and Rb phosphorylation.**

Asynchronous HCT116, MCF7, and RPE-1 hTERT cells were treated with CDK4/6 inhibitors (CDK4/6-in) or vehicle control for up to 48 hours. DNA content was analysed by flow cytometry, and phosphorylated Rb at serine 780 (pRb) levels were examined by Western blot. **a**, Schematic representation of the experimental time-course setup. **b**, Flow cytometry analysis of cell cycle profiles (G1, S, G2/M) in HCT116 cells treated with increasing concentrations of palbociclib (Palbo). **c**, Representative immunoblot showing pRb, total Rb, and β-tubulin (β-Tub) levels in HCT116 cells treated with Palbo for 24 hours (top). Quantification of pRb band intensities normalized to total Rb is plotted, with $IC_{50}$ and $C_{max}$ values indicated (bottom). **d, e**, HCT116 cells were treated with increasing concentrations of Palbo, abemaciclib (Abema), ribociclib (Ribo), or the MEK inhibitor PD0325901 (MEKi) for 24 hours. **d**, Flow cytometry analysis of cell cycle profiles. **e**, Western blot analysis of pRb, total Rb, and β-tubulin levels. Quantified pRb levels normalized to total Rb are shown in bar graphs. **f**, MCF7 cells treated with Palbo were analysed by flow cytometry for cell cycle profiles at 24 hours. **g, h**, RPE-1 hTERT cells treated with Palbo were analysed. **g**, Flow cytometry analysis of DNA content. **h**, Western blot analysis of pRb, total Rb, and β-tubulin. Quantified pRb levels are plotted with $IC_{50}$ and $C_{max}$ values estimated.

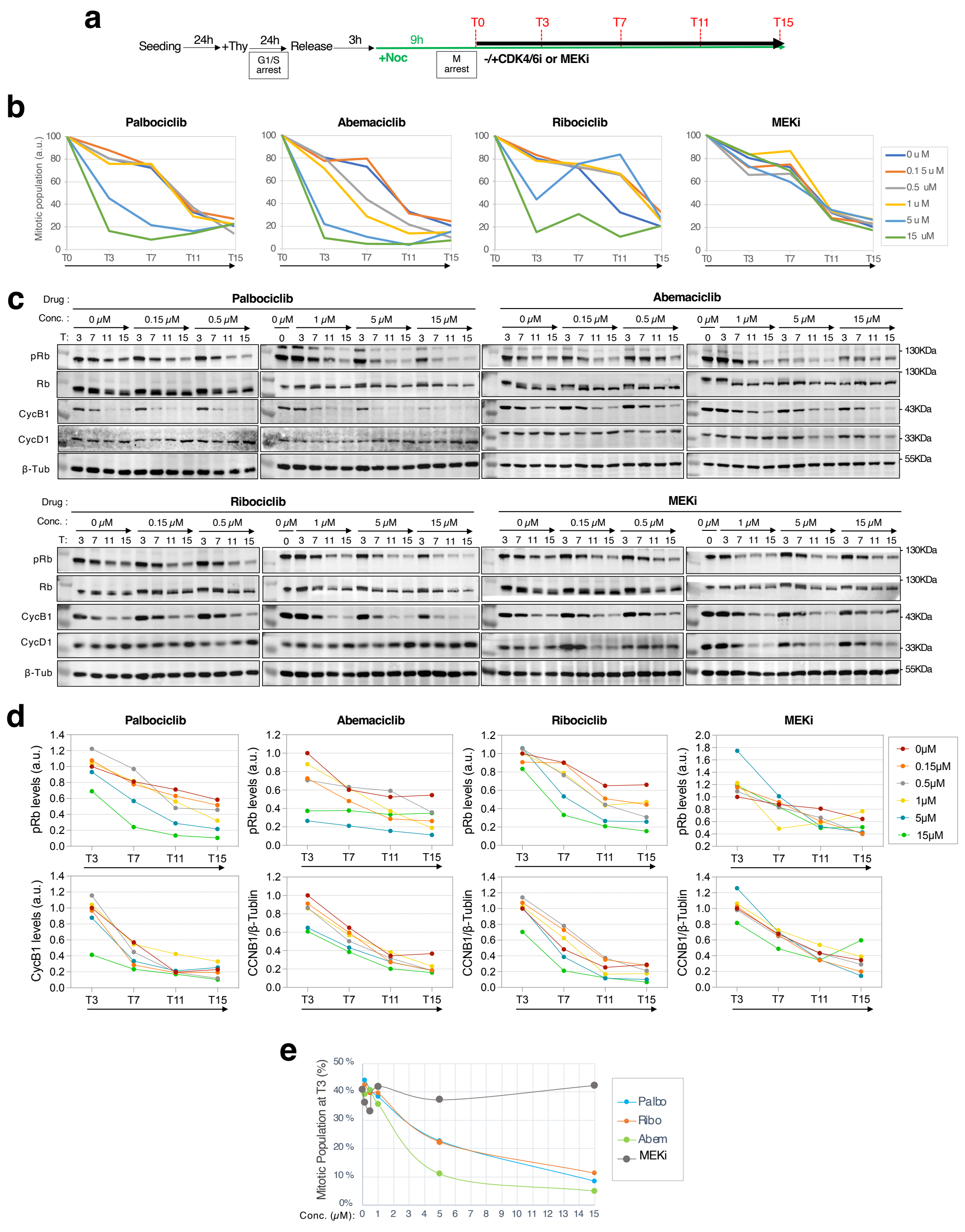


**Extended Data Figure 2 CDK4/6 inhibitors induce premature mitotic exit in nocodazole-arrested cells**

**Extended Data Figure 2 CDK4/6 inhibitors induce premature mitotic exit in nocodazole-arrested cells.**

**a,** Experimental scheme: HCT116 cells were synchronised at the G1/S boundary, released into nocodazole-containing medium for 9 hours to arrest in prometaphase, and treated with CDK4/6 inhibitors or MEK inhibitor (MEKi). Samples were collected at the indicated time points for analysis. **b,** Relative mitotic populations (4N DNA content, pH3-positive) following treatment with increasing concentrations of palbociclib, abemaciclib, ribociclib, or MEKi were quantified by flow cytometry and plotted as arbitrary units, with the mitotic population at T0 set as 100. CDK4/6 inhibitors induced a dose-dependent reduction in mitotic cells, with abemaciclib being the most potent. **c,** Representative Western blot images showing phosphorylated Rb (pRb), total Rb, Cyclin B1 (CycB1), Cyclin D1 (CycD1), and β-tubulin (loading control) levels after treatment with CDK4/6 inhibitors or MEKi. **d,** Quantification of pRb and CycB1 band intensities from Western blot analyses. Relative levels at each time point were normalised to loading controls and plotted as arbitrary units. **e,** Comparative kinetics of mitotic exit induced by CDK4/6 inhibitors and MEKi based on mitotic populations at T3 (3 hours after addition). Abemaciclib induced the fastest mitotic exit, followed by palbociclib and ribociclib, while MEKi showed no significant effect.

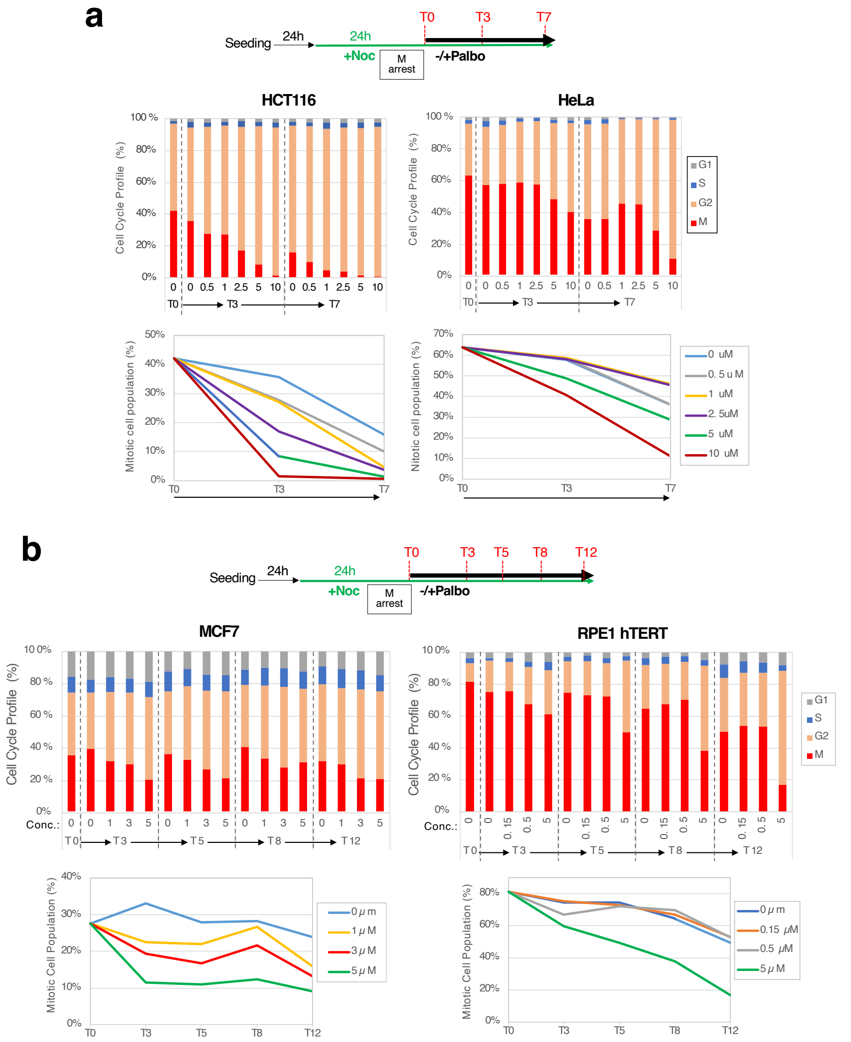


**Extended Data Figure 3 CDK4/6 inhibition induces premature mitotic exit in diverse cell lines**

**Extended Data Figure 3 CDK4/6 inhibition induces premature mitotic exit in diverse cell lines.**

**a, b,** Mitotic slippage following CDK4/6 inhibition was assessed in HCT116, HeLa, MCF7, and RPE-1 hTERT cells. Cells were treated with nocodazole for 24 hours to induce mitotic arrest (4N DNA content with pH3 signals). Palbo was added at the indicated concentrations, and samples were collected at specified time points for flow cytometry analysis. **a,** HCT116 and HeLa cells were treated with Palbo at 0, 0.5, 1, 2.5, 5, or 10 µM, and harvested at 0, 3, and 7 hours post-treatment. Line graphs show the proportion of mitotic cells (4N DNA content with positive pH3 signals). **b,** MCF7 and RPE-1 hTERT cells were treated with Palbo at 0, 1, 3, or 5 µM, and harvested at 0, 3, 5, 8, and 12 hours post-treatment. Similar to HCT116 cells, Palbo induced accelerated mitotic slippage in all cell lines, although the sensitivity to Palbo varied among the lines.

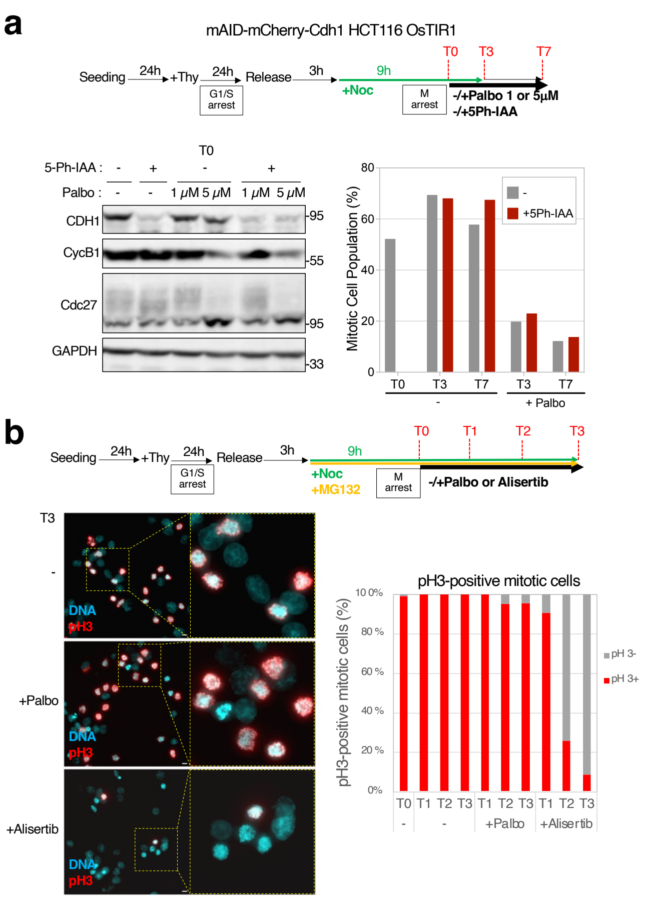


**Extended Data Figure 4 CDK4/6 inhibition-induced premature mitotic exit is independent of APC/C$^{CDH1}$ and Aurora B kinase**

**Extended Data Figure 4 CDK4/6 inhibition-induced premature mitotic exit is independent of APC/CCDH1 activity and Aurora B kinase.**

**a,** Premature mitotic exit upon CDK4/6 inhibition is not mediated by $APC/C^{CDH1}$. HCT116 cells expressing OsTIR1 and mAID-mCherry-tagged CDH1 were synchronised in prometaphase with nocodazole treatment. Depletion of CDH1 was induced by the addition of 5-Ph-IAA (500 μM), reducing CDH1 levels to below 20% after 3 hours (T0). Palbo was added at T0, and samples were collected at the indicated time points. Flow cytometry and Western blot analysis showed that mitotic exit and CycB1 degradation occurred with comparable kinetics in CDH1-depleted and control cells, indicating that CDH1 is not required for mitotic exit induced by CDK4/6 inhibition. **b,** Aurora B inhibition by alisertib triggers mitotic exit with loss of pH3 signals, while CDK4/6 inhibition does not affect pH3 levels in mitotic cells. HCT116 cells were treated with Palbo or alisertib after nocodazole-induced mitotic arrest. Immunofluorescence staining revealed that alisertib reduced pH3 signals despite the presence of condensed mitotic chromosomes, whereas Palbo-treated cells retained pH3 signals. Scale bar: 10 μm. Quantification of pH3-positive and -negative mitotic cells is shown as a bar graph.

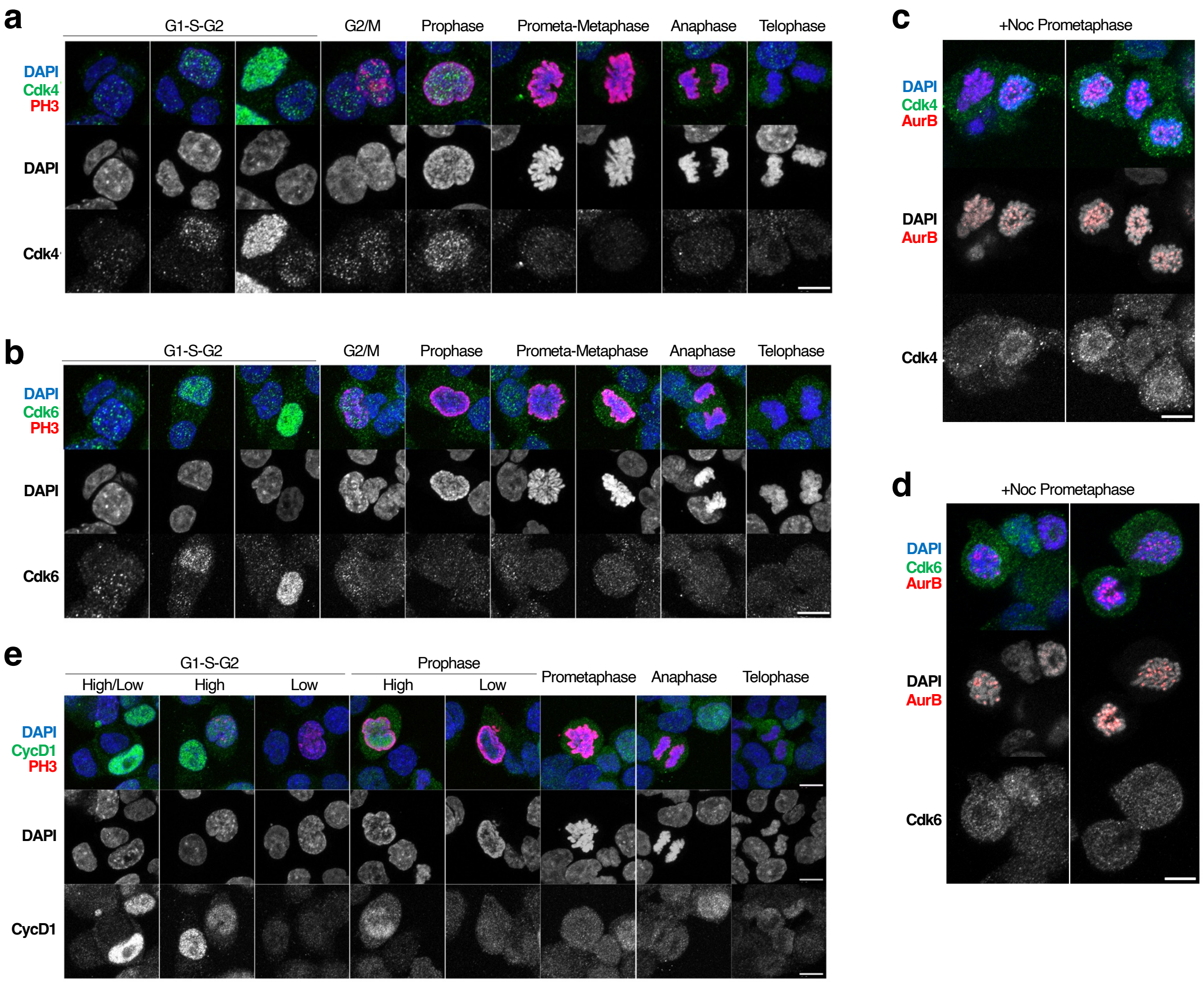


**Extended Data Figure 5 CDK4 is associated with mitotic chromosomes**

**Extended Data Figure 5 CDK4 associates with chromosomes during early mitosis and SAC arrest.**

**a, b,** Subcellular localisation of CDK4 and CDK6 was examined in asynchronous HCT116 cells using immunofluorescence staining. CDK4 and CDK6 predominantly localised to the nucleus during interphase and were dispersed as cells progressed through mitosis. CDK4 exhibited a weak association with chromosomes during early mitosis (prophase to prometaphase). Chromosome morphology and pH3 staining were used to determine cell cycle phases. Scale bar: 10 µm. **c, d,** Localisation of CDK4 during SAC arrest. HCT116 cells were synchronised at G1/S by a thymidine block, released into nocodazole-containing medium, and harvested after 9 hours to arrest in prometaphase. Immunofluorescence staining revealed pronounced CDK4 association with chromosomes during SAC-dependent mitotic arrest. Aurora kinase B (AurB) served as an inner kinetochore marker. Scale bar: 10 µm. **e,** Cyclin D1 (CycD1) showed nuclear localisation in interphase and weak association with chromosomes during early mitosis (prophase to prometaphase). CycD1 expression levels varied between cells, reflecting fluctuations in mitogenic signals. pH3 staining indicated mitotic cells. Scale bar: 10 µm.
